# Experimental metal contamination reduces gut microbiota diversity and alters its composition and function in wild-caught fish

**DOI:** 10.1101/2025.03.21.644596

**Authors:** Quentin Petitjean, Séverine Jean, Margarita Granada, Sophie Manzi, Charlotte Veyssiere, Annie Perrault, Myriam Cousseau, Pascal Laffaille, Lisa Jacquin, Joël White

**Author notes:** These authors contributed equally to this work (co-senior authors) (Lisa Jacquin, Joël White). Corresponding author: Quentin PETITJEAN.

## Abstract

Wild organisms face environmental stressors that can interact and affect their health unexpectedly. Evidence suggests that responses to stressors may be mediated by changes in the gut microbiota, with cascading effects on host health. However, the combined effects of multiple stressors on host microbiota are still overlooked. Here, we investigated the single and interactive effects of realistic metal contamination (i.e., a mixture of Cd, Cu, Zn) and an immune challenge (i.e., lipopolysaccharides - LPS and phytohaemagglutinin - PHA) mimicking a parasite attack on the taxonomic and functional diversity and composition of gut microbiota among several wild freshwater fish (*Gobio occitaniae*) populations sampled along a gradient of contamination in streams. We found that the experimental metal contamination strongly altered the gut microbial community, with no interaction with the immune secondary stressor. Indeed, metal contamination reduced both taxonomic and functional gut microbial diversity, affecting the microbial community’s taxonomic and functional composition, with predicted consequences for their functional role in fish. Metal contamination reduced microbial function related to molecule biosyntheses (e.g., cell structure and amino acid precursors) while increasing functions associated with energy production (e.g., anaerobic respiration). In addition, populations sampled along a gradient of pollution in the wild did not differ in their response, suggesting a consistent impact of contaminants irrespective of the host’s past exposure to pollution. Our results highlight how realistic levels of metal contamination alter the fish gut microbiota, potentially affecting their ability to cope with environmental stressors, though long-term fitness implications are still unclear.

## 1 Introduction

Human activities alter biodiversity and ecosystem functioning by causing numerous disturbances [1]. Stressors rarely occur in isolation in a natural setting, and it is often difficult to predict their combined effects on biodiversity, especially in aquatic systems [2]. To tackle this challenge, many studies have investigated how stressors may interact and subsequently alter the traits of freshwater organisms (e.g., [3–5]). However, few studies have explored how multiple stressors alter host-associated microbiota (but see [6–8]), especially in fish, despite their key role in host physiology, immune responses, development, and behavior [9–13].

Understanding the effects of environmental stressors on the gut microbiota is crucial for assessing the host’s ability to acclimate or adapt to contrasting environmental conditions [14, 15]. While genetic adaptation and phenotypic plasticity play an essential role in the evolutionary responses to stress [16], the microbiota may also modulate host phenotypes, potentially affecting fitness and evolutionary trajectories [17–19]. For instance, gut microbiota can detoxify toxic compounds, as shown in wood rats (*Neotoma* spp.) [20]. Xenobiotics such as insecticides or antibiotics may interact with gut microbes, detoxifying or producing secondary metabolites with deleterious effects on host health [21, 22]. Host genotype and environment shape gut microbiota and, in turn, affect host phenotypic traits and plasticity [19]. Therefore, it is essential to study microbiota variations among populations that have experienced different stress levels.

Fish are well-suited models for studying gut microbiota variability [12, 23, 24], as their gut microbial communities vary with genotype [25], ecology—especially diet [9, 24]—and environmental stressors [13, 26], though contaminants effects remain overlooked [27]. Some studies suggest metal contamination can alter fish gut microbiota and impact host health [28–30]; for example, cadmium exposure in Nile tilapia (*Oreochromis niloticus*) reduced microbial diversity and community dissimilarity [29]. In contrast, pathogen-induced microbiota changes are better documented, especially in aquaculture [31]. Several studies link disease severity to gut microbiome disruption (e.g., [32–34]); for instance, largemouth bronze gudgeon (*Coreius guichenoti*) infected by furunculosis (*Aeromonas salmonicida*) showed reduced α-diversity and altered β-diversity with decreased community dissimilarity and dispersion compared to healthy individuals [35]. Metal contamination and pathogen infections may interact in complex ways to shape fish microbiota. Indeed, metals can impair immune function, particularly inflammatory responses [4, 36], resulting in cascading multistress effects on fish gut microbiota. However, the combined impact of metals and pathogens remains unclear due to limited experimental studies conducted in wild, heterogeneous fish populations.

In this study, we investigated the effect of an environmentally relevant mixture of metals most frequently found in French streams (i.e., Cadmium – Cd, Copper – Cu, and Zinc – Zn) and/or of an immune challenge (i.e., antigen mix mimicking parasite attack) on the gut microbiota of a benthic freshwater fish (*Gobio occitaniae)* under controlled conditions. To assess intraspecific variation, we compared gut microbiota responses across five populations from low- and high-pollution environments [37].

According to previous studies, we first expect a decrease in gut microbial α-diversity (e.g., richness) in fish exposed to metal contamination or immune stress, as stress can strongly impact microbial communities [38]. We also expect such patterns to be altered by combined exposure to both stressors (i.e., antagonistic or synergistic effect), reflecting multistress effects.

Second, we expect significant shifts in community composition in response to metal contamination [39], immune challenge [31], and/or their combination depending on the population’s origin (i.e., legacy effect). Indeed, chronic exposure to pollutants, as shown in sediment microbial communities, can induce legacy effects that dampen responses to additional stressors [40].

Third, stressors may affect microbiota β-diversity in opposing ways. Stressors may reduce interindividual variability (i.e., homogenization) through deterministic processes selecting for more similar, low diversity, dysbiotic microbiota [29, 35, 41]. Alternatively, stressors may increase interindividual variability through stochastic processes (i.e., Anna Karenina Principle—AKP) [42], as seen in mice under enzymatic stress, likely due to opportunistic taxa invasion facilitated by a compromised immune system [43]. Both patterns have been reported depending on stressor type and host species [44], though some fish studies support a homogenizing effect of metal and immune stressors on gut microbial communities [29, 35].

Fourth, we hypothesize that single and combined stressors would induce dysbiosis (i.e., changes in microbial ratio reflecting a dysfunctional community) as documented in earlier studies [45–47]. We expect an increase in opportunistic or potentially pathogenic taxa alongside a decline in beneficial ones. This includes changes in the relative abundance of *Firmicutes*, *Bacteroidetes*, and *Proteobacteria*, affecting F/P and B/P ratios commonly associated with dysbiosis [39].

Finally, we predict that stressors would reduce functional α-diversity by altering community diversity and composition, leading to the potential loss of non-redundant functions. We also expected shifts in functional β-diversity, as stressors have been shown to impact various metabolic pathways [48, 49]. While effects on functional heterogeneity (i.e., interindividual variance) remain uncertain, we anticipated changes under metal exposure—especially in pathways related to oxidative stress and energy metabolism [48, 49]—though not necessarily under the immune challenge alone.

## 2 Material and Methods

### 2.1 Model Species and Sampling Sites

The gudgeon (Gobio occitaniae) is a benthic freshwater fish species from Southwestern France characterized by low mobility [50, 51], thus chronically exposed to metal contamination via water and sediments in human-altered streams. Based on previous field surveys [52, 53], fish populations were collected from three low-pollution (FARM, ARIMAS, CELCAB) and two high-pollution (AUSCOR, RIOU) areas, mainly contaminated with Cd, Cu, and Zn (Table 1, see also [4] for site details and electrofishing methods). Fish were then transported to the laboratory, treated with praziquantel (Vetofish, France; purity: 1000 mg.g⁻¹; 3 mg.L⁻¹) to eliminate parasites and standardize immune status, and acclimated under controlled laboratory conditions for 30 days ([4] for details).

**Table 1.** Experimental design and final sample sizes of fish gut collected at the end of the experiment. NC: Non-contaminated. C: Contaminated. LP: Low Pollution. HP: High Pollution. PBS: control injection of Phosphate-Buffered Saline. AMIX: Injection with a mixture of antigens. (See [4] for more details.)

| Population | Lat. | Long. | Fish origin | NC-PBS | NC-AMIX | C-PBS | C-AMIX | TOTAL |
| --- | --- | --- | --- | --- | --- | --- | --- | --- |
| FARM | 2°20'9.1255" E | 45°38'50.0269" N | Low Pollution (LP) | 13 | 10 | 15 | 11 | 49 |
| ARIMAS | 1°22'26.1494" E | 43°5'2.4677" N | Low Pollution (LP) | 11 | 14 | 14 | 14 | 53 |
| CELCAB | 1°39'27.8050" E | 44°30'27.5836" N | Low Pollution (LP) | 8 | 3 | 12 | 6 | 29 |
| AUSCOR | 1°19'38.2922" E | 43°39'11.3598" N | High Pollution (HP) | 28 | 26 | 29 | 38 | 121 |
| RIOU | 2°12'41.3680" E | 44°33'55.4580" N | High Pollution (HP) | 13 | 15 | 15 | 12 | 55 |
| TOTAL |  |  |  | 73 | 68 | 85 | 81 | 307 |

### 2.2 Experimental Design

Fish were experimentally exposed in a full factorial design to environmentally realistic metal contamination (C; based on concentrations from the highly polluted site RIOU: Cd, 14 µg.LLJ¹; Cu, 10 µg.LLJ¹; Zn, 600 µg.LLJ¹) and/or immune challenge (AMIX), mimicking parasite attack [54]. The AMIX treatment consisted of phytohaemagglutinin-P (Sigma-Aldrich, USA; 90 µg.10 µLLJ¹) and *Escherichia coli* lipopolysaccharide (serotype O111:B4, Sigma-Aldrich, USA; 90 µg.10 µLLJ¹) according to a previous study conducted in gudgeons [54]. Control injection treatment consisted of a neutral Phosphate Buffered Saline (PBS) solution. The four experimental treatment groups were: NC-PBS (no contamination, PBS injection), C-PBS (contamination, PBS), NC-AMIX (no contamination, antigen mixture), and C-AMIX (contamination, antigen mixture) (see Table 1 and [4] for details).

Along experimental exposure, fish were housed in groups of five in tanks (50×30×30 cm) at constant temperature (17–18 °C) with a 12:12 h photoperiod and daily feeding (JBL Propond Sterlet S pellets; ∼1% biomass). Fish were randomly exposed to control or contaminated water for 14 days, with PBS or AMIX injection administered on day 7. Initially, 410 fish were introduced, showing no significant differences in sex ratio (χ^2^ = 2.53, p = 0.47), mass (8.64 ± 4.77g; Anova: F = 0.21, p = 0.89), or size (9.4 ± 1.7cm; Anova: F = 0.57, p = 0.63) among treatment [4]. After exposure, 313 gut samples were collected; 6 were excluded due to poor DNA, leaving 307 gut samples for analysis (Table 1). Tank water samples were also collected to characterize microbial communities in water.

### 2.3 DNA Extraction, PCR Amplification, And 16s rRNA Sequencing

Fish guts were stored at −20°C before DNA extraction using DNeasy® Blood & Tissue Kits (Qiagen, Netherlands) under UV-treated laminar flow conditions. Briefly, gut samples were cut into pieces and lysed (40 min at 37°C) with TE buffer Tris-EDTA 10X, Triton™ X-100, nuclease-free water (Fisher Scientific, USA), and Lysozyme (Dutscher, France), followed by proteinase K digestion within AL lysis buffer (30 min, 50°C). DNA purification was performed through ethanol precipitation and successive column washes (DNeasy® mini spin column, buffers AW1, AW2), with final elution in AE/C buffer according to manufacturer recommendations (Qiagen, Netherlands). DNA concentration and purity were verified using a NanoDrop 1000 (Thermo-Fisher Scientific, USA).

PCR amplification of the microbial 16S rRNA V5-V6 region used primers 5′-GGATTAGATACCCTGGTAGT-3′ and 5′-CACGACACGAGCTGACG-3′ in 20 µL reactions containing AmpliTaq Gold™ 360 Master Mix (10µL, Fisher Scientific, USA), primers (1 μL at 0.4 µM each), Bovine Serum Albumin (0.16μL, Sigma-Aldrich, USA), nuclease-free water (5.84μL) and 2 µL of undiluted DNA extract. PCR steps included initial denaturation (95°C, 10 min), 30 cycles of denaturation (95°C, 30 s), hybridization (57°C, 30 s), elongation (72°C, 90 s), and a final elongation (72°C, 7 min).

Sequencing was performed in triplicate, pooling 4μL of amplicons per sample (one library per replicate), at the Genotoul-Bioinfo platform (Toulouse, France) using Illumina MiSeq technology (2 × 300 bp paired-end V3 chemistry, Illumina, USA).

Extraction, PCR, and sequencing negative controls, along with ZymoBIOMICS™ Microbial Community DNA Standard (ZYMO Research, USA) positive control, were included in the design to control for potential contamination, PCR effectiveness, and tag-jump. All reagents used were molecular biology grade.

### 2.4 Data Analysis

#### 2.4.1 Bioinformatic Pipeline

Bioinformatic analyses of raw 16S DNA sequences were performed on the genologin bioinformatic server hosted at the <u>Genotoul-Bioinfo platform</u> (Toulouse, France) using OBITools package [55]. Paired-end reads (R1 and R2) were assembled, demultiplexed, and filtered to exclude sequences with alignment scores <50, non-attributed nucleotides (Ns), or length <50 bp. Duplicates and singletons were removed, sequencing triplicates merged, and sequences clustered into Molecular Operational Taxonomic Units (MOTUs) at 97% similarity using SUMACLUST v1.0.31 [56]. Taxonomic annotations were performed using the SILVAngs reference database for small sequences (16S and 18S; v138.1, August 2020) [57].

#### 2.4.2 Data Pre-Processing

MOTUs with relative abundance <0.005% or present in fewer than three samples were removed following procedures reported in [58]. Four samples without remaining reads were discarded. The metabaR R package [59] was used to further remove artifacts and contaminants and evaluate sequencing depth, PCR replicability, and lower tag-jump effects. MOTUs were classified as contaminants if their frequency in negative controls exceeded that in samples, resulting in the removal of six MOTUs (6 MOTUs out of 487, 1% of total MOTUs; Supplementary Material 1) and 0.4% of reads (25,051 out of 5,169,994). Although sequencing depth was <1000 reads in approximately 55% of samples (including tank water), PCR replicability was high (95%). After quality filtering, one additional sample was discarded (Table 1).

#### 2.4.3 Statistical Analyses

All analyses except taxonomic annotation and pathway inference (Bash scripts, Ubuntu 22.04.3 LTS) were performed using R v4.2.3. Data and code to replicate the analyses are available at the following URLs:Data: <u>10.5281/zenodo.14989874</u>; Code: <u>10.5281/zenodo.14990006</u>.

##### 2.4.3.1 Normalization Strategy

Since normalization methods can affect interpretations of 16S DNA sequence data [e.g., 13– 17], we evaluated the consistency of α- and β-diversity results across five common normalization methods (i.e., proportion, cumulative sum scaling - CSS, rarefaction, DESeq, and trimmed mean of M-value - TMM) and their log-transformed (log2(x+1)) equivalent. Results were generally consistent across methods (Supplementary Material 2); thus, we present only findings based on the original log-transformed dataset for clarity.

##### 2.4.3.2 α-Diversity

We first explored the effects of experimental stressors (metal contamination and immune challenge) on gut microbial α-diversity using taxonomic and phylogenetic indices computed with Vegan [65] and Picante [66] R packages. Specifically, we calculated MOTU richness, Shannon index and its exponential, Simpson index, and Faith’s phylogenetic diversity (see Supplementary Material 3 for details about phylogenetic tree constructed using DECIPHER [67] and phangorn [68] R packages).

Effects of contamination (NC vs. C), immune challenge (PBS vs. AMIX), pollution level at fish origin (Low vs. High), and their third-order interaction were tested using linear mixed models (LMMs; lme4 R package [69]). Fish size and sex were included as covariates, with the experimental tank nested within the session (trial), and the identity of the population of origin as random effects. The best models were selected based on the lowest corrected Akaike’s Information Criterion (AICc) using the MuMIn package [70]. When models differed by ΔAICc < 2, the model with the highest weight was retained.

##### 2.4.3.3 β-Diversity

Second, we assessed the effects of stressors on gut microbial community composition (β-diversity) using multiple distance metrics: Bray-Curtis (abundance-based), Jaccard (presence/absence), Robust Aitchison (relative abundance), Hellinger (major MOTUs emphasis), UniFrac and Weighted UniFrac (presence/absence and abundance, respectively accounting for phylogenetic distance), computed with Vegan [65] and phyloseq [71] R packages.

We evaluated correlations between experimental treatments, covariates (i.e., fish sex and size), and ordination scores from Principal Coordinate Analysis (PCoA) using envfit (Vegan R package [65]; detailed in Supplementary Material 7). Then, scores from the two primary PCoA axes were analyzed using linear mixed models (LMMs), with the best models selected having the lowest AICc (as previously described). Finally, group dispersion homogeneity (beta dispersion) was assessed using betadisper (Vegan package [65]) to compare within-group variances among treatments significantly affecting β-diversity.

##### 2.4.3.4 Functional Annotation

Functional annotation (i.e., pathway inference) of microbial MOTUs was performed using PICRUSt2 software [72] to infer potential microbial community functions from MOTU sequences. Briefly, MOTU sequences were formatted into a Biological Observation Matrix (BIOM) [73], followed by sequence placement and hidden state prediction for 16S DNA copy. MOTUs with weighted Nearest Sequenced Taxon Index (NSTI) scores >2 were removed (one MOTU excluded). The average NSTI score across samples was low (0.07 ± 0.003 SE), suggesting reliable predictions of microbial functions. We then performed metagenome predictions for enzyme classification and inferred pathway abundance based on the presence of gene families according to the Metacyc database [74].

##### 2.4.3.5 Taxonomic and Functional Analyses

To identify microbial taxa and functions affected by stressors, we performed differential abundance (DA) analyses using linear mixed models (LMMs) implemented within the LinDA framework (Microbiotatat R package [75]). Analyses were conducted at the phylum and family levels for taxa and primarily at the function class and superpathway levels for functional pathways, excluding taxa and functions detected in fewer than five samples.

We further assessed microbial community alteration using two ratios considered as indicators of dysbiosis in the literature: *Firmicutes/Bacteroidetes* (F/B) and *Proteobacteria/Bacteroidetes* (P/B) ratios (e.g., [45–47]), analyzed via LMMs. Model structures testing the effects of stressors on both dysbiosis ratio and DA were similar to those previously described for α-diversity analyses, but here, we used backward selection to retain significant terms. P-values were adjusted using the false discovery rate method [76].

## 3 Results

### 3.1 α-Diversity

Experimental metal contamination significantly decreased all gut α-diversity indices (except Simpson’s index) for MOTUs and inferred functions, irrespective of fish origin or immune challenge (Figure 1A, Table 2).

**Figure 1.**
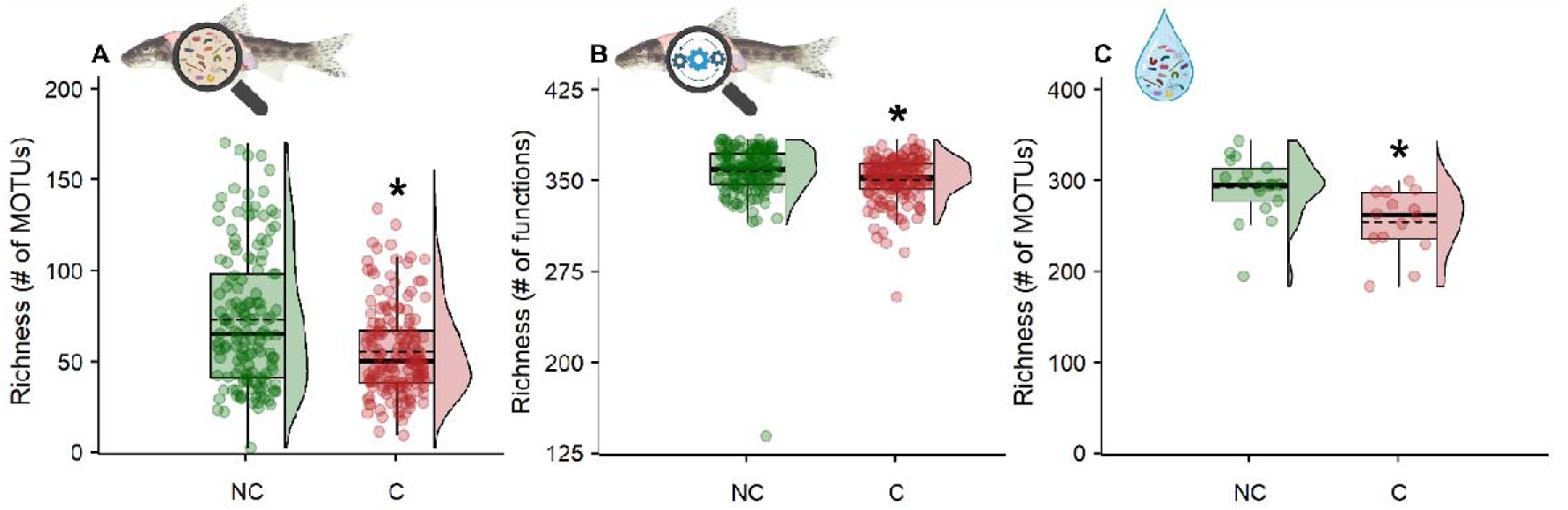
(A) MOTUs and inferred functions richness in fish gut (A and B respectively), and MOTUs richness in water (B) according to the experimental metal contamination treatment (Green: non-contaminated control; red: contaminated treatment). Stars above boxplots indicate significant differences (LMM, p-value < 0.05) between contamination treatments. The density distribution is represented on the right side of each boxplot.

**Table 2.** Effects of experimental metal contamination (NC vs. C), immune challenge (PBS vs. AMIX), and the fish origin (LP vs. HP) on α-diversity indices (Richness, exp(Shannon), Simpson and Faith’s PD) on fish gut microbiota community (MOTUs, left) and inferred functional composition (from picrust2, right) performed on original log-transformed data. Results are obtained from the best (selected using AICc) final linear mixed models (LMM). Sample size and marginal and conditional R square are reported as n, R2m, and R2c, respectively. Faith’s PD index was not computed for functional composition as it relies on phylogenetic distances among taxa.

| Taxonomic diversity |  |  |  |  |  |  | Functional diversity |  |  |  |  |  |
| --- | --- | --- | --- | --- | --- | --- | --- | --- | --- | --- | --- | --- |
|  | Estimate | Std. Error | t value | df | Chisq | p.value | Estimate | Std. Error | t value | df | Chisq | p.value |
| **Richness \| n = 307 \| R2m = 0.177 \| R2c = 0.302** |  |  |  |  |  |  | **Richness \| n = 307 \| R2m = 0.14 \| R2c = 0.24** |  |  |  |  |  |
| Intercept | 101 | 12.9 | 7.8 | 1 | 60.8 | <0.0001 | 387 | 8.77 | 44.1 | 1 | 1940 | <0.0001 |
| Contamination (NC) | 18.2 | 3.52 | 5.17 | 1 | 26.7 | <0.0001 | 7.22 | 2.29 | 3.16 | 1 | 9.96 | <0.01 |
| Sex (M) | 15.7 | 3.83 | 4.1 | 2 | 17.4 | <0.01 | 8.71 | 2.61 | 3.34 | 2 | 11.2 | <0.01 |
| Size (cm) | -5.15 | 1.18 | -4.38 | 1 | 19.1 | <0.0001 | -3.91 | 0.799 | -4.89 | 1 | 23.9 | <0.0001 |
| **Shannon.exp \| n = 307 \| R2m = 0.172 \| R2c = 0.298** |  |  |  |  |  |  | **Shannon.exp \| n = 307 \| R2m = 0.16 \| R2c = 0.282** |  |  |  |  |  |
| Intercept | 80 | 10 | 7.98 | 1 | 63.7 | <0.0001 | 368 | 9.11 | 40.4 | 1 | 1630 | <0.0001 |
| Contamination (NC) | 13.6 | 2.7 | 5.03 | 1 | 25.3 | <0.0001 | 8.12 | 2.34 | 3.47 | 1 | 12 | <0.01 |
| Sex (M) | 12.2 | 2.96 | 4.12 | 2 | 17.7 | <0.01 | 8.91 | 2.67 | 3.34 | 2 | 11.2 | <0.01 |
| Size (cm) | -3.95 | 0.91 | -4.34 | 1 | 18.8 | <0.0001 | -4.51 | 0.821 | -5.49 | 1 | 30.2 | <0.0001 |
| **Simpson \| n = 307 \| R2m = 0.0654 \| R2c = 0.113** |  |  |  |  |  |  | **Simpson \| n = 307 \| R2m = 0.0869 \| R2c = 0.16** |  |  |  |  |  |
| Intercept | 0.991 | 0.00931 | 106 | 1 | 11300 | <0.0001 | 0.997 | 0.000135 | 7370 | 1 | 54400000 | <0.0001 |
| Contamination (NC) | 0.00374 | 0.00257 | 1.46 | 1 | 2.13 | 0.145 | 0.0000682 | 0.0000365 | 1.87 | 1 | 3.49 | 0.0617 |
| Sex (M) | 0.0078 | 0.00292 | 2.67 | 2 | 7.2 | <0.05 | 0.000101 | 0.0000415 | 2.43 | 2 | 5.92 | 0.0518 |
| Size (cm) | -0.00266 | 0.000884 | -3.01 | 1 | 9.06 | <0.01 | -0.0000493 | 0.0000127 | -3.88 | 1 | 15.1 | <0.01 |
| **PD \| n = 307 \| R2m = 0.138 \| R2c = 0.256** |  |  |  |  |  |  |  |  |  |  |  |  |
| Intercept | 16.9 | 1.72 | 9.87 | 1 | 97.4 | <0.0001 |  |  |  |  |  |  |
| Contamination (NC) | 1.7 | 0.447 | 3.81 | 1 | 14.5 | <0.01 |  |  |  |  |  |  |
| Sex (M) | 1.96 | 0.508 | 3.86 | 2 | 15.9 | <0.01 |  |  |  |  |  |  |
| Size (cm) | -0.654 | 0.156 | -4.18 | 1 | 17.5 | <0.0001 |  |  |  |  |  |  |

Specifically, fish exposed to contaminated water (C) exhibited lower MOTU richness (56.8±3.1) compared to uncontaminated controls (NC: 73.8±1.9; Figure 1A). Similarly, functional richness decreased under contamination (C: 351.9±1.4; NC: 358.8±2.0; Figure 1B). In contrast, immune challenge (PBS vs. AMIX) and fish origin (LP vs. HP) had no significant effects (fixed factors removed from final model, Table 2). Fish sex and size significantly affected α-diversity, with males and juveniles having higher diversity indices than females and smaller fish exhibiting greater MOTU and functional diversity (Table 2).

Metal contamination significantly reduced microbial α-diversity in water samples, decreasing MOTU richness from 293.7 ± 8.2 (NC) to 256.1 ± 9.4 (C; Figure 1C, Supplementary Material 4).

### 3.2 β-Diversity

Experimental metal contamination significantly affected β-diversity indices (MOTUs and inferred functions) in fish gut microbial communities, irrespective of fish origin or immune challenge (Figure 2, Table 3; Supplementary Material 2 for comparisons across normalization methods).

**Figure 2.**
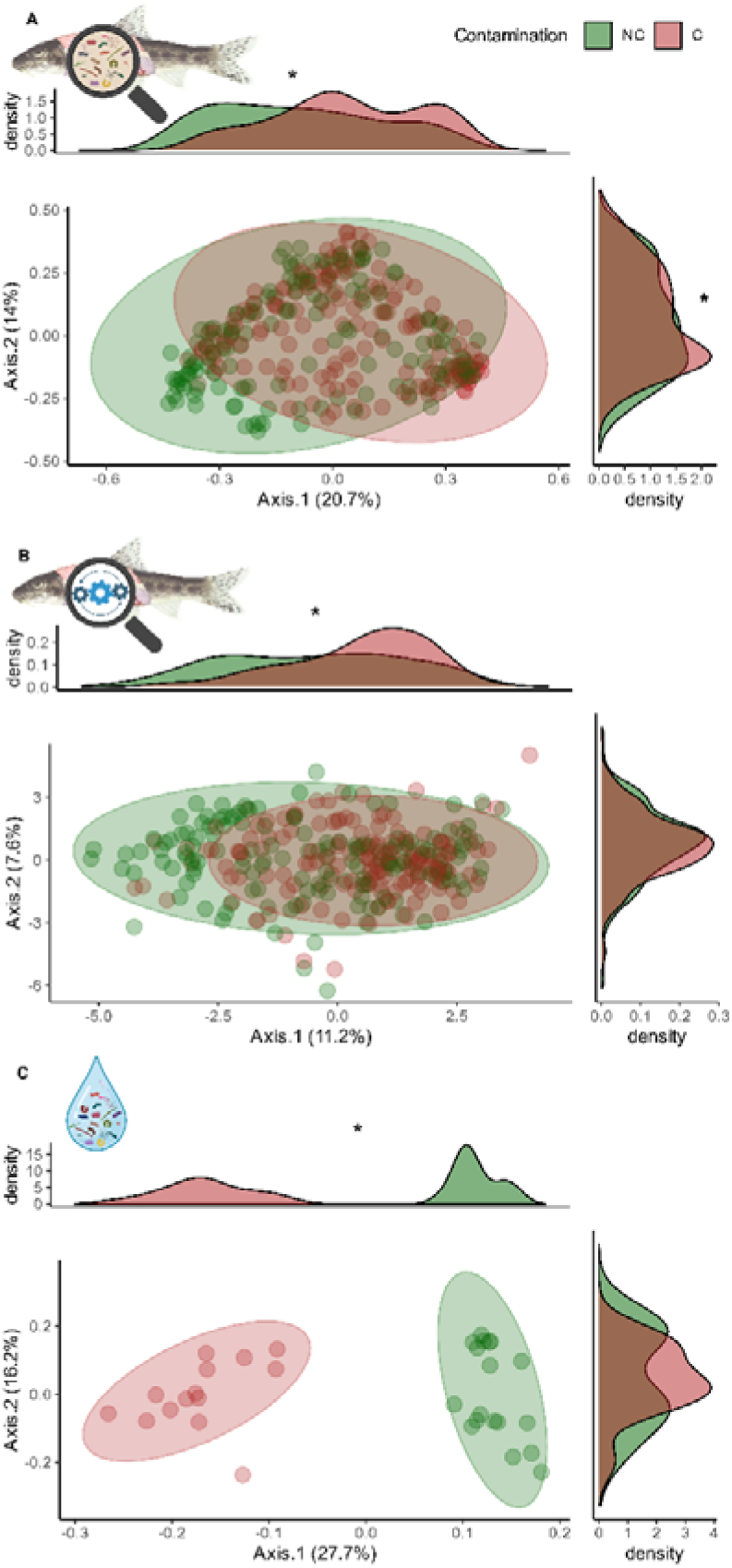
Dissimilarity of microbial community (MOTUs) and inferred functions in fish gut (A and B respectively) and dissimilarity of microbial community in water (C). A and C display PCOA performed on Bray-Curtis distances, while B displays PCOA performed on Robust Aitchison distances according to the experimental metal contamination treatment (Green: non-contaminated control; red: contaminated treatment). Density distribution along axis 1 and 2 are represented above and on the right of each PCOA axis, respectively.

**Table 3.** Effects of experimental metal contamination (NC vs. C), immune challenge (PBS vs. AMIX), fish origin (NC vs. C) and covariates (fish size and sex) on PCOA scores (first axis) representing β-diversity indices (Bray Curtis, Jaccard, Robust Aitchison, Hellinger, Weighed Unifrac and Unweighted Unifrac) on fish gut microbiota community (MOTUs, left) and inferred functional composition (from picrust2, right) performed on original log-transformed data. Results are obtained from the best (selected using AICc) final linear mixed models (LMM). Sample size and marginal and conditional R square are reported as n, R2m, and R2c, respectively. Unifrac and weighed Unifrac distances were not computed for functional composition as they rely on phylogenetic distances among taxa.

| Taxonomic composition |  |  |  |  |  |  | Functional composition |  |  |  |  |  |
| --- | --- | --- | --- | --- | --- | --- | --- | --- | --- | --- | --- | --- |
|  | Estimate | Std. Error | t value | df | Chisq | p.value | Estimate | Std. Error | t value | df | Chisq | p.value |
| **Bray \| n = 307 \| R2m = 0.0934 \| R2c = 0.557** |  |  |  |  |  |  | **Bray \| n = 307 \| R2m = 0.162 \| R2c = 0.387** |  |  |  |  |  |
| Intercept | -0.189 | 0.0871 | -2.17 | 1 | 4.73 | <0.05 | -0.209 | 0.0506 | -4.12 | 1 | 17 | <0.0001 |
| Contamination (NC) | -0.104 | 0.0208 | -5.01 | 1 | 25.1 | <0.0001 | -0.0544 | 0.013 | -4.2 | 1 | 17.6 | <0.0001 |
| Sex (M) | -0.0644 | 0.0215 | -3 | 2 | 8.98 | <0.05 | -0.0451 | 0.0142 | -3.18 | 2 | 10.3 | <0.01 |
| Size (cm) | 0.0239 | 0.00658 | 3.62 | 1 | 13.1 | <0.01 | 0.0247 | 0.00437 | 5.66 | 1 | 32 | <0.0001 |
| **Jaccard \| n = 307 \| R2m = 0.0912 \| R2c = 0.566** |  |  |  |  |  |  | **Jaccard \| n = 307 \| R2m = 0.152 \| R2c = 0.386** |  |  |  |  |  |
| Intercept | -0.162 | 0.0792 | -2.05 | 1 | 4.21 | <0.05 | -0.28 | 0.0719 | -3.9 | 1 | 15.2 | <0.0001 |
| Contamination (NC) | -0.097 | 0.0187 | -5.18 | 1 | 26.8 | <0.0001 | -0.0786 | 0.0185 | -4.25 | 1 | 18.1 | <0.0001 |
| Sex (M) | -0.0565 | 0.0194 | -2.92 | 2 | 8.5 | <0.05 | -0.0599 | 0.0201 | -2.98 | 2 | 9 | <0.05 |
| Size (cm) | 0.0207 | 0.00593 | 3.5 | 1 | 12.2 | <0.01 | 0.0334 | 0.0062 | 5.39 | 1 | 29 | <0.0001 |
| **Robust.Aitchison \| n = 307 \| R2m = 0.136 \| R2c = 0.343** |  |  |  |  |  |  | **Robust.Aitchison \| n = 307 \| R2m = 0.148 \| R2c = 0.443** |  |  |  |  |  |
| Intercept | -1.3 | 0.633 | -2.06 | 1 | 4.24 | <0.05 | -2.79 | 0.777 | -3.6 | 1 | 12.9 | <0.01 |
| Contamination (NC) | -0.961 | 0.186 | -5.18 | 1 | 26.8 | <0.0001 | -0.989 | 0.192 | -5.15 | 1 | 26.5 | <0.0001 |
| Sex (M) | -0.508 | 0.187 | -2.72 | 2 | 7.41 | <0.05 | -0.576 | 0.208 | -2.76 | 2 | 7.71 | <0.05 |
| Size (cm) | 0.189 | 0.0572 | 3.3 | 1 | 10.9 | <0.01 | 0.334 | 0.0643 | 5.19 | 1 | 27 | <0.0001 |
| **Hellinger \| n = 307 \| R2m = 0.0555 \| R2c = 0.717** |  |  |  |  |  |  | **Hellinger \| n = 307 \| R2m = 0.167 \| R2c = 0.281** |  |  |  |  |  |
| Intercept | -0.17 | 0.111 | -1.54 | 1 | 2.36 | 0.125 | -0.0923 | 0.032 | -2.89 | 1 | 8.33 | <0.01 |
| Contamination (NC) | -0.114 | 0.0239 | -4.76 | 1 | 22.6 | <0.0001 | -0.0427 | 0.00846 | -5.05 | 1 | 25.5 | <0.0001 |
| Sex (M) | -0.0588 | 0.0226 | -2.6 | 2 | 6.84 | <0.05 | -0.037 | 0.00948 | -3.9 | 2 | 15.3 | <0.01 |
| Size (cm) | 0.0204 | 0.00691 | 2.95 | 1 | 8.7 | <0.01 | 0.0126 | 0.00291 | 4.34 | 1 | 18.8 | <0.0001 |
| **WUniFrac \| n = 307 \| R2m = 0.105 \| R2c = 0.239** |  |  |  |  |  |  |  |  |  |  |  |  |
| Intercept | -0.0227 | 0.032 | -0.709 | 1 | 0.503 | 0.478 |  |  |  |  |  |  |
| Contamination (NC) | -0.0448 | 0.00931 | -4.82 | 1 | 23.2 | <0.0001 |  |  |  |  |  |  |
| Sex (M) | -0.0311 | 0.00994 | -3.13 | 2 | 10.3 | <0.01 |  |  |  |  |  |  |
| Size (cm) | 0.00517 | 0.00299 | 1.73 | 1 | 2.99 | 0.0837 |  |  |  |  |  |  |
| **UniFrac \| n = 307 \| R2m = 0.133 \| R2c = 0.219** |  |  |  |  |  |  |  |  |  |  |  |  |
| Intercept | -0.168 | 0.0694 | -2.42 | 1 | 5.86 | <0.05 |  |  |  |  |  |  |
| Contamination (NC) | -0.0824 | 0.0186 | -4.43 | 1 | 19.6 | <0.0001 |  |  |  |  |  |  |
| Sex (M) | -0.0748 | 0.0211 | -3.54 | 2 | 13.3 | <0.01 |  |  |  |  |  |  |
| Size (cm) | 0.0233 | 0.00647 | 3.6 | 1 | 13 | <0.01 |  |  |  |  |  |  |

Specifically, fish exposed to contamination (C) differed significantly from uncontaminated controls (NC) in terms of gut microbial community composition (Figure 2A, Table 3; Supplementary Material 5 for PCOA axis 2) and inferred functions (Figure 2B, Table 3; Supplementary Material 5). Neither immune challenge nor fish origin significantly affected the microbial community and inferred functions (Table 3; Supplementary Material 5). However, fish sex and size significantly affected microbial composition and inferred functions (Table 3; Supplementary Material 5).

Metal contamination and covariates (sex, size) had mostly no effect on group dispersion homogeneity (BetaDisper approach; Supplementary Material 8). However, focusing on inferred functions in fish gut, we found that fish from HP sites exhibited higher functional heterogeneity than fish from LP sites (fish origin) according to Bray and Jaccard indices, regardless of the contamination treatment. (Supplementary Material 8).

Finally, metal contamination significantly altered microbial community composition in water samples, especially along the first ordination axis (Figure 2C; Supplementary Material 6).

### 3.3 Differential Taxonomic Abundance

Experimental metal contamination significantly affected DA of gut MOTUs, irrespective of fish origin (Figure 3; Supplementary Materials 9 and 10). At the family level (Figure 3A; Supplementary Material 9), contamination significantly decreased the abundance of several taxa, primarily low-abundance families such as *Enterococcaceae, Devosiaceae, Microbacteriaceae*, but also the highly abundant *Aeromonadaceae*. At the phylum level, contamination decreased abundances of *Proteobacteria*—the most abundant phylum—and *Actinobacteriota* and *Planctomycetota* (Figure 3B; Supplementary Material 10).

**Figure 3.**
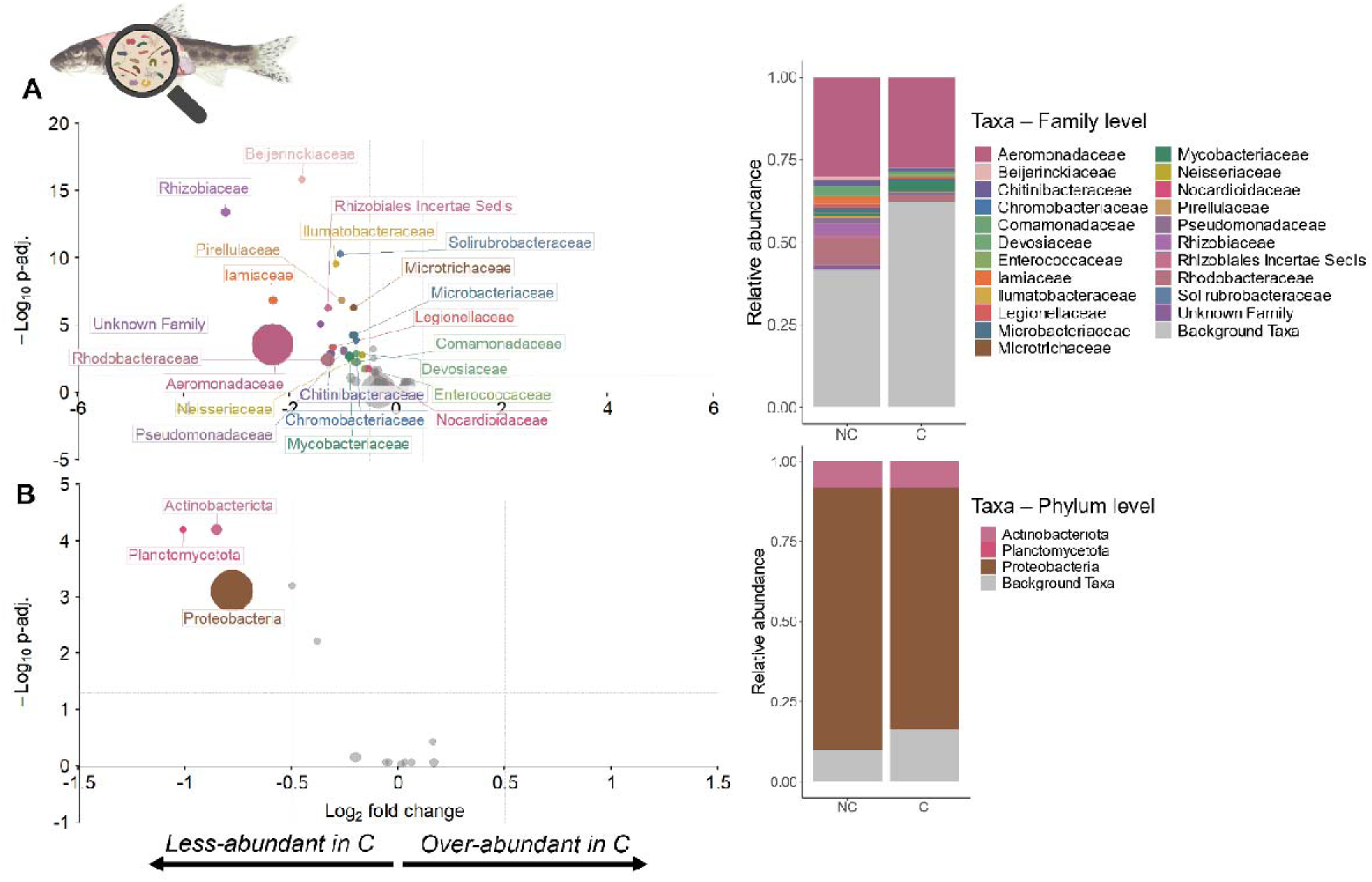
Differential abundance (DA) of MOTUs according to the experimental metal contamination treatment identified by LINDA. The left panel (A and B) shows the relative abundances of microbial taxa in the gut of fish exposed to contaminated (C) vs. non-contaminated (NC) water, shown at the family level (A) and phylum level (B) according to a p-value threshold of 0.05 and a log2FC above |0.5|. The size of the dots is proportional to the number of read counts. The right panel (A and B) shows the relative abundance of the significantly affected taxa and background (non-affected) taxa at the family (A) or phylum (B) level. Grey indicates no significant differential abundance, whereas colored markers represent significantly less- or over-abundant taxa.

Immune challenge (PBS vs. AMIX) had limited effects, significantly increasing only *Pseudomonadaceae* abundance at the family level (Supplementary Material 9), while fish origin (LP vs. HP) had no significant effects at either taxonomic level. Also, F/B and P/B ratios remained unaffected by stressors (Supplementary Material 11).

In water samples, metal contamination significantly affected DA of MOTUs at both family and phylum levels. Specifically, contamination decreased the abundance of several families, such as *Aeromonadaceae*, *Microbacteriaceae*, and *Microtrichaceae*, and increased others, such as *Flavobacteriaceae*, *Chitinophagaceae*, and *Neisseriaceae* (Supplementary Material 12). Similarly, at the phylum level, *Actinobacteriota*, *Proteobacteria*, and *Firmicutes* decreased, while *Bacteroidota* increased (Supplementary Material 13). Notably, While metal contamination similarly affected some taxa (e.g., *Aeromonadaceae*, *Microbacteriaceae*) in both gut and water, other taxa (e.g., *Neisseriaceae*, *Bacteroidota*) responded differently depending on the environment (Supplementary Materials 9, 10, 12, and 13). Additionally, Rhodobacteraceae were more abundant in tanks housing fish from HP sites (Supplementary Material 12).

### 3.4 Differential Functional Abundance

Experimental metal contamination significantly affected the DA of MetaCyc pathways in fish guts, irrespective of fish origin (Figure 4; Supplementary Material 14). Specifically, contamination decreased predicted abundance of pathways involved in cell structure biosynthesis, secondary metabolites (i.e., organic compounds not directly involved in growth, development, and reproduction), and demethylmenaquinone (a molecule involved in electron transfer in bacteria membrane), as well as C1 compounds (carbon oxidation to CO^2^) and amino acid metabolism (Figure 4, Supplementary Material 15). Conversely, contamination increased pathways linked to anaerobic respiration and sugar degradation (degradation of sugars to provide energy; Figure 4, Supplementary Material 15). However, altered pathways showed low predicted abundances (<1%; Figure 4B); thus, these results should be taken with caution.

**Figure 4.**
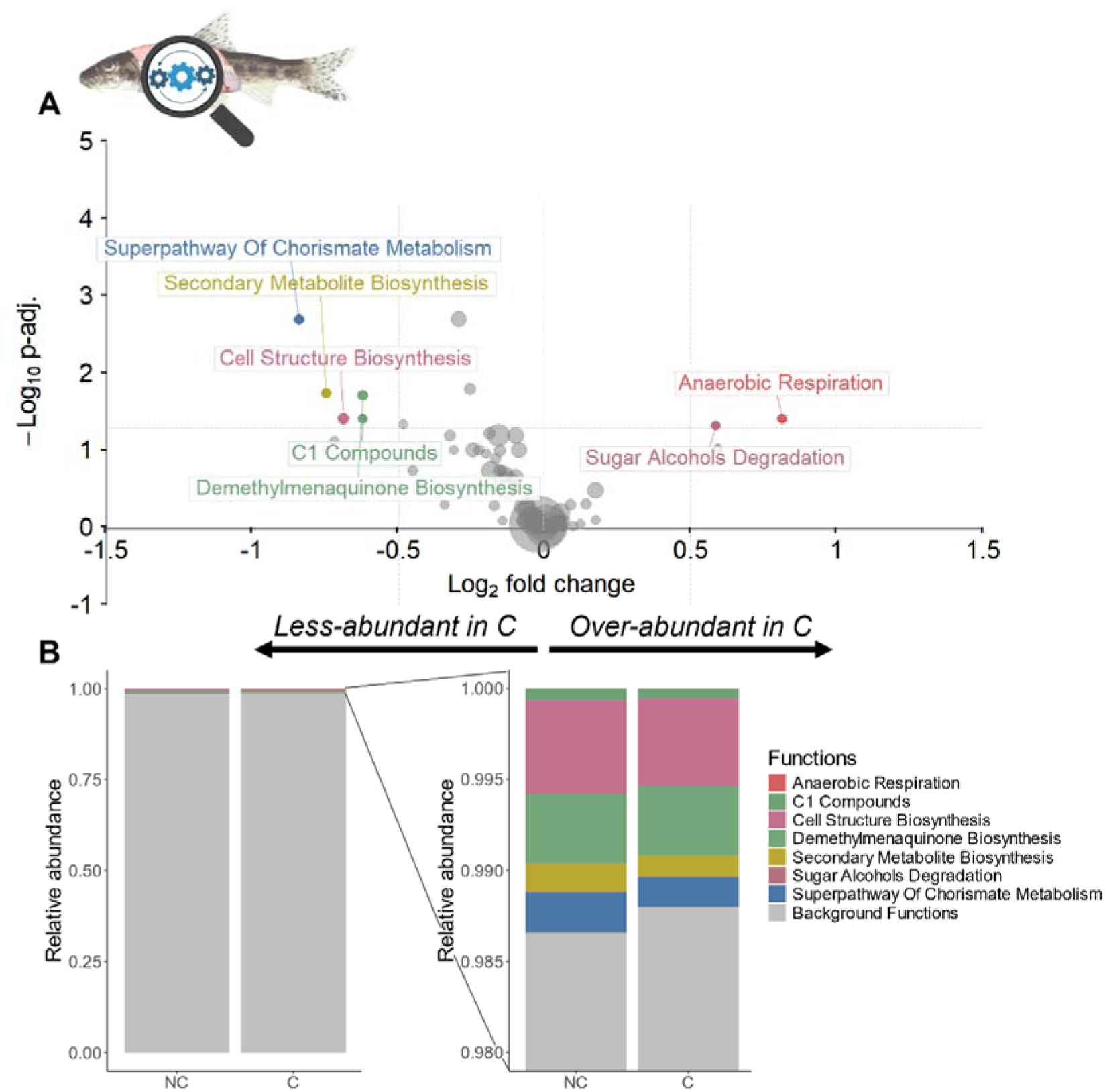
Differential abundance of inferred functions according to the experimental metal contamination treatment identified by LINDA. A) shows the relative abundances of inferred functions (picrust2 and Metacyc pathways) in the gut of fish exposed to contaminated (C) vs. non-contaminated (NC) water, according to a p-value threshold of 0.05 and a log2FC above |0.5|. The size of the dots is proportional to the abundance of functions. B) Display the relative abundance of the significantly affected and background (non-affected) functions according contaminated (C) vs. non-contaminated (NC) treatments. The right panel corresponds to a zoom on the upper part of the right barplot. Grey indicates no significant differential abundance, whereas colored markers represent significantly less- or over-abundant taxa.

Immune challenge (PBS vs. AMIX) and fish origin (LP vs. HP) did not significantly affect the DA of inferred functions in fish gut (Supplementary Material 12).

## 4 Discussion

Taken together, our results show that metal contamination strongly affected fish gut microbial communities in wild gudgeons (*Gobio occitaniae*) with little or no effects of immune challenge or fish origin. We also found no interactive effect between metal exposure and immune challenge on the gut microbiota.

More specifically, the experimental metal contamination reduced α-diversity and altered fish gut microbiota’s community and functional composition (β-diversity). However, stressors did not alter gut microbial communities dispersion (i.e., interindividual variance). Rather, contamination seems to lead to deterministic and similar changes in microbial gut communities under metal pollution, whatever the additional stressors and the origin of the fish population. The experimental contamination mostly reduced the relative abundance of low-abundance families and *Actinobacteriota*, *Planctomycetota*, and *Proteobacteria* phylum, with no clear effect on commonly used dysbiosis ratios. Such alteration of the microbial community was accompanied by a decline in predicted functions related to cell structure biosynthesis, electron transfer, carbon oxidation, and amino acid and secondary metabolite production. In contrast, metal contamination increased the predicted abundance of functions tied to anaerobic respiration and sugar alcohol degradation. Altogether, it suggests that immune challenge and intraspecific variability (i.e., population origin) have little effect on fish gut microbiota, whereas metal contamination of the water shapes fish gut microbiota and alters microbial functions, potentially explaining some changes in fish health found in previous studies as discussed below.

### 4.1 Effects of Contaminant on microbial diversity

Exposure to environmentally realistic metal contamination (Cd, Cu, Zn) under controlled conditions significantly reduced gut microbial diversity (i.e., Richness, Shannon index, Faith’s PD) in Gudgeons. This aligns with previous studies linking metal exposure [23, 77, 78]—and contaminants more broadly [39, 79, 80]—to altered gut microbial diversity, though the direction of reported effects may vary.

Surprisingly, metal contamination did not significantly affect the Simpson index, suggesting contamination impacts less abundant MOTUs more strongly (i.e., the Simpson index emphasizes dominant species). Since low-abundance taxa can play key functional roles [81], the observed effect may further affect community dissimilarity.

The concurrent decline in diversity in fish gut and surrounding contaminated water supports the “biodiversity hypothesis,” which posits that reduced environmental microbial diversity can lead to gut microbial deprivation and impaired host health [82–84]. Alternatively, metal contamination may independently shape gut and water microbiota by similarly selecting for or against specific taxa, as suggested by the result of the DA analysis (e.g., A*tinobacteriota* and *Proteobacteria* phylum).

### 4.2 Contamination effects on microbial composition

Overall, we found that over 98% of bacteria in wild gudgeon guts belonged to four dominant phyla: *Proteobacteria (79%), Firmicutes (8.1%), Actinobacteriota (8.0%),* and *Bacteroidota (3.6%).* This composition aligns with previous studies reporting *Proteobacteria, Firmicutes*, *Bacteroides*, and *Actinobacteriota* as fish’s most common gut taxa [85, 86].

Experimental contamination significantly altered the community composition in both fish gut and water. More particularly, metal contamination decreased the relative abundance of MOTUs belonging to 22 families and 3 phyla in fish gut and 49 families and 7 phyla in water. Most MOTUs with altered DA in the gut of fish exposed to contamination were relatively low abundant except those belonging to the *Aeromonadaceae* family and *Proteobacteria* phylum, supporting the results emphasized by the Simpson index (see previous section). Interestingly, some taxa, such as *Aeromonadaceae*, *Microbacteriaceae and Microtrichaceae* (family level), and *Actinobacteriota* and *Proteobacteria* (phylum level), exhibited parallel decreases in DA in both fish gut and water, suggesting a direct influence of environmental microbial changes on the gut microbiota.

Nonetheless, the shifts in water microbial communities alone cannot fully account for the changes in the gut community of fish exposed to experimental contamination, suggesting that host filtering might also play a role. For instance, the relative abundance of *Neisseriaceae* (family level) increased in water but declined in the gut, and *Bacteroidota* (phylum level) rose in water without a corresponding gut increase. Overall, 11 families (50% of taxa with altered DA in the gut) and two phyla (75% of taxa with altered DA in the gut) showed consistent declines in both gut and water. While underlying mechanisms remain unclear, the effect of host filtering on the gut microbial community has previously been proposed to be related to genotype, immunity (see [87] for review), and epigenetic feedback [88]. In a previous study on the same fish populations, metal exposure induced phenotypic shifts in immunity and behavior [4]. These findings highlight the complex interplay between environmental microbiota and host regulatory mechanisms in shaping gut microbial responses to stressors. Whatever the underlying mechanisms, our findings align with previous studies reporting altered relative abundances for various microbial taxa in the gut of fish exposed to metal contamination (see [39] for review). Unlike previous studies, the experimental metal contamination did not increase the relative abundance of any MOTUs in fish guts. Indeed, other studies have observed taxonomic shifts, such as a rise in *Firmicutes* and a decline in *Proteobacteria* in zebrafish (*Danio rerio*) exposed to waterborne lead (Pb) [30] or an increase in *Bacteroides* and *Proteobacteria* alongside a decrease in *Fusobacteria* in Nile tilapia exposed to waterborne Cd [29]. In addition, our results show that contamination did not affect commonly used dysbiosis indicators (F/B and P/B ratios) despite shifts in microbial composition and abundance. The finding highlights the potential limitations of relying on these ratios as indicators of microbial community alterations in fish guts under contaminated conditions.

Our study adds evidence to the growing body of research suggesting that some microbial taxa may be favored or disfavored under contaminants exposure, depending on the host species, the contaminant types, and, more broadly, the environmental conditions [17]. This highlights the need for further research to identify patterns and predict their effects on gut microbiota function and host health.

### 4.3 Contamination effects on microbial functions

Beyond taxonomic shifts, we found that experimental metal contamination reduced functional diversity and altered the functional composition of the gut microbiota. Specifically, contamination decreased functions related to biosynthesis—cell structure, secondary metabolites, aromatic amino acid precursors (e.g., tryptophan, tyrosine, phenylalanine), and mediators of electron transfer—while increasing functions tied to energy production, particularly anaerobic respiration and sugar alcohol degradation. Notably, tryptophan, tyrosine, and phenylalanine are essential amino acids that animals cannot synthesize and must obtain from their diet [89]. In fish, these are synthesized by bacteria through chorismate metabolism, a pathway that is also less abundant in the guts of fish exposed to metal contamination. Tryptophan and tyrosine, especially, are involved in forming neurotransmitters such as serotonin and dopamine [90], potentially affecting immune responses [91], social and feeding behavior [92, 93], and cognition [94]. In a previous study, we observed metal-induced changes in fish immune (i.e., neutrophil/lymphocyte ratio) and behavioral traits (e.g., swimming, exploration) [4]. These findings raise important questions about whether gut microbiota functional shifts contribute to such physiological and behavioral changes, with broader implications for wild fish health.

In contrast, metal contamination increased microbial functions related to energy production *via* anaerobic respiration and degradation of sugar derived from carbohydrates and starches, potentially supplying carbon and energy to the host. This suggests that metal contamination may promote anaerobic conditions in the gut and reshape microbial communities, though the mechanisms remain unclear. Accordingly, we observed reduced abundances of aerotolerant taxa such as *Proteobacteria* and *Actinobacteria* [95] in the gut of metal-exposed fish. Further studies are now needed to understand the mechanisms and consequences of such changes for the hosts.

Overall, our results suggest that metal contamination primarily affected fish gut microbiota by reducing diversity, mainly through a decrease in the relative abundance of low-abundance taxa. This alteration of the microbial community reduced the abundance of some inferred functions in fish guts, with potential consequences on fish traits such as immune responses, behavior, and ability to mobilize energy. However, the functions altered were of low relative abundance (<1%), raising questions about their biological relevance for host health. A key limitation comes from using inference methods to predict the functions provided by the microbial community in the fish gut; thus, future metagenomic and metatranscriptomic approaches are needed to clarify the functional impacts of metal—and broader stressor-induced—microbial changes.

### 4.4 Transient Effects of Immune Challenge and Limited Influence of Fish Origin

Contrary to our expectations, we found very few effects of either immune challenge, combined metal contamination and immune challenge, and fish origin (LP—Low pollution vs. High Pollution—HP) on the gut microbial community. This is surprising given the known interplay between immunity and host microbiota [11, 12]. We previously showed that the antigen mixture used here effectively triggered local and systemic immune responses with physiological effects, confirming its efficacy at the tested dose [54]. One possibility is that the immune challenge caused transient microbial shifts that recovered within the 7-day window before sampling. In line with this idea, we observed an increased relative abundance of *Pseudomonadaceae* in immune-challenged fish, suggesting a short-term microbial response that may have partly recovered [96]. However, a longitudinal study is needed to test this hypothesis. Whatever the underlying mechanism, the immune challenge did not alter the microbiota’s response to metal exposure, likely due to the strong dominant effect of metal contamination found in a previous study on fish traits [4].

Surprisingly, we found no effect of fish origin on the gut microbiota, contradicting our hypothesis of high intraspecific variability linked to contrasting native environments (i.e., legacy effect [40]). This result might reflect the homogenizing influence of standardized laboratory conditions and commercial diets throughout acclimatization and experimentation, thereby revealing the strong effect of the experimental metal contamination of the water. Consistently with previous studies [97], any initial differences in gut microbiota among wild populations might have been hindered by captivity. Interestingly, the relative abundance of *Rhodobacteraceae* increased in water from tanks housing fish from polluted sites, possibly due to differences in mucus-associated microbiota, as this family is commonly found in fish mucus [98]. This suggests that differences in mucus microbiota according to fish origin could alter the microbial community in the surrounding environment (i.e., experimental unit) and potentially bias experimental outcomes under laboratory conditions. Accounting for fish origin is thus crucial when studying microbiota–stress interactions. Further studies in the field, for instance, using a reciprocal transplant experimental design [37], will be valuable in exploring microbiota divergence in more realistic field conditions and under natural diets [99].

### 4.5 Anna Karenina Principle, pollution-induced deterministic changes or stochasticity?

The AKP predicts that hosts not exposed to stressors share similar “healthy” microbiota (due to deterministic and predictable changes), while stressed individuals exhibit divergent microbial communities (i.e., stochastic dispersion within groups) [42]. Contrary to this hypothesis, our results showed no increase in within-group dispersion under stress, instead supporting a consistent, deterministic effect of stressors on gut microbiota across fish populations. This aligns with our prediction and the results of previous field studies [41, 100] and a meta-analysis [96], suggesting that disturbed microbiota are not always more variable [42] but may adopt a new (predictable) deterministic configuration under pollution exposure, whatever the fish origin. These findings indicate that we may be able to predict shifts in microbial communities in contaminated environments. However, further empirical studies across a wider range of environmental conditions and wild species are necessary to generalize this hypothesis.

## 5 Conclusion

Taken together, our findings highlight that realistic metal contamination found in streams significantly alters the gut microbiota in a wild fish, mostly by reducing microbial diversity, especially through the loss of low-abundance taxa, and altering community structure and the functions provided by bacteria. Indeed, changes in the microbial community in fish guts reduce microbial functions that may, in turn, affect the host’s energy mobilization, immune responses, and behavior. Also, our results suggest that metal contamination does not drive change in the microbial community in a stochastic way but rather in a deterministic one, contradicting the Anna Karenina principle. Finally, this effect of metal contamination was also consistent whatever the origin of fish populations and their past exposure in the wild (Low Pollution or High Pollution origin). Although the long-term implications for fitness remain to be investigated, our results suggest that fish gut microbiota is altered by environmental contaminants, affecting non-redundant microbial functions and potentially impairing key physiological processes essential for fish health. Further studies are needed to investigate environmental stressors’ effects on gut-associated bacteria under field-realistic conditions to decipher the effect of intrinsic and extrinsic factors in shaping the gut microbial communities and infer the potential consequences on host health.

## Acknowledgments

The authors warmly thank the CRBE laboratory, notably the AQUAECO and ECI teams. They also thank the members of the GeT-PlaGe platform, Amaia IRIBAR, and Uxue SUESCUN, for their help in DNA sequencing.

## 6 Declarations

### 6.3 Compliance With Ethical Standards

Experimental procedures complied with French and European legislation for animal experimentation (European directive 2010/63/UE) and were conducted under the French animal handler’s certificate (N°31-103), the establishment approval for vertebrate experimentation N°A3113002, and were approved by the ethical committee n°073 (authorization n°8538). Fish sampling was conducted under local authorities’ authorization, and fish were treated for parasites according to the prescription N°2529 delivered by VetoFish.

### 6.4 Availability of Data and Materials

The data and Rscripts supporting this study’s findings are openly available in the following zenodo repositories: Data: 10.5281/zenodo.14989874; Code: 10.5281/zenodo.14990006.

### 6.5 Authors’ Contributions

LJ, SJ, JW, PL and QP conceived the ideas and designed the methodology; LJ, SJ, CV, SM, PL, AP, MC and QP conducted the experiments and/or collected the data; MG and QP analyzed the data; QP led the writing of the manuscript. All authors contributed critically to the drafts.

## References

1. Reid AJ et al. Emerging threats and persistent conservation challenges for freshwater biodiversity. Biological Reviews 2019;94:849–873. 10.1111/brv.12480

2. Petitjean Q et al. Stress responses in fish: From molecular to evolutionary processes. Science of The Total Environment 2019;684:371–380. 10.1016/j.scitotenv.2019.05.357

3. Marcogliese DJ, Pietrock M. Combined effects of parasites and contaminants on animal health: parasites do matter. Trends in Parasitology 2011;27:123–130. 10.1016/j.pt.2010.11.002

4. Petitjean Q et al. Intraspecific variability of responses to combined metal contamination and immune challenge among wild fish populations. Environmental Pollution 2020;116042. 10.1016/j.envpol.2020.116042

5. Jacquin L et al. High temperature aggravates the effects of pesticides in goldfish. Ecotoxicology and Environmental Safety 2019;172:255–264. 10.1016/j.ecoenv.2019.01.085

6. Maher RL et al. Multiple stressors interact primarily through antagonism to drive changes in the coral microbiome. Sci Rep 2019;9:6834. 10.1038/s41598-019-43274-8

7. Zaneveld JR et al. Overfishing and nutrient pollution interact with temperature to disrupt coral reefs down to microbial scales. Nat Commun 2016;7:11833. 10.1038/ncomms11833

8. Li JD et al. Dual stressors of infection and warming can destabilize host microbiomes. Philosophical Transactions of the Royal Society B: Biological Sciences 2024;379:20230069. 10.1098/rstb.2023.0069

9. Colston TJ, Jackson CR. Microbiome evolution along divergent branches of the vertebrate tree of life: what is known and unknown. Mol Ecol 2016;25:3776–3800. 10.1111/mec.13730

10. Hsiao EY et al. Microbiota Modulate Behavioral and Physiological Abnormalities Associated with Neurodevelopmental Disorders. Cell 2013;155:1451–1463. 10.1016/j.cell.2013.11.024

11. Rea K, Dinan TG, Cryan JF. The microbiome: A key regulator of stress and neuroinflammation. Neurobiology of Stress 2016;4:23–33. 10.1016/j.ynstr.2016.03.001

12. Diwan AD, Harke SN, Panche AN. Host-microbiome interaction in fish and shellfish: An overview. Fish and Shellfish Immunology Reports 2023;4:100091. 10.1016/j.fsirep.2023.100091

13. Butt RL, Volkoff H. Gut Microbiota and Energy Homeostasis in Fish. Front Endocrinol 2019;10. 10.3389/fendo.2019.00009

14. Alberdi A et al. Do Vertebrate Gut Metagenomes Confer Rapid Ecological Adaptation? Trends in Ecology & Evolution 2016;31:689–699. 10.1016/j.tree.2016.06.008

15. Decaestecker E et al. Hierarchical eco-evo dynamics mediated by the gut microbiome. Trends in Ecology & Evolution 2024;39:165–174. 10.1016/j.tree.2023.09.013

16. Hendry AP. Eco-evolutionary Dynamics. Princeton University Press, 2016.

17. Henry LP et al. The microbiome extends host evolutionary potential. Nat Commun 2021;12:5141. 10.1038/s41467-021-25315-x

18. McKenney EA et al. The ecosystem services of animal microbiomes. Mol Ecol 2018;27:2164– 2172. 10.1111/mec.14532

19. Lange C et al. Impact of intraspecific variation in insect microbiomes on host phenotype and evolution. ISME J 2023;17:1798–1807. 10.1038/s41396-023-01500-2

20. Kohl KD, Dearing MD. The Woodrat Gut Microbiota as an Experimental System for Understanding Microbial Metabolism of Dietary Toxins. Front Microbiol 2016;7. 10.3389/fmicb.2016.01165

21. Peterson BF. Microbiome toxicology — bacterial activation and detoxification of insecticidal compounds. Current Opinion in Insect Science 2024;63:101192. 10.1016/j.cois.2024.101192

22. Collins SL, Patterson AD. The gut microbiome: an orchestrator of xenobiotic metabolism. Acta Pharm Sin B 2020;10:19–32. 10.1016/j.apsb.2019.12.001

23. Suzzi AL et al. Legacy metal contamination is reflected in the fish gut microbiome in an urbanised estuary. Environmental Pollution 2022;314:120222. 10.1016/j.envpol.2022.120222

24. Clements KD et al. Intestinal microbiota in fishes: what’s known and what’s not. Molecular Ecology 2014;23:1891–1898. 10.1111/mec.12699

25. Bolnick DI et al. Major Histocompatibility Complex class IIb polymorphism influences gut microbiota composition and diversity. Molecular Ecology 2014;23:4831–4845. 10.1111/mec.12846

26. Wang AR et al. Progress in fish gastrointestinal microbiota research. Reviews in Aquaculture 2018;10:626–640. 10.1111/raq.12191

27. Adamovsky O et al. The gut microbiome and aquatic toxicology: An emerging concept for environmental health. Environmental Toxicology and Chemistry 2018;37:2758–2775. 10.1002/etc.4249

28. Meng X-L et al. Intestinal microbiota and lipid metabolism responses in the common carp (*Cyprinus carpio* L.) following copper exposure. Ecotoxicology and Environmental Safety 2018;160:257–264. 10.1016/j.ecoenv.2018.05.050

29. Zhai Q et al. Effect of dietary probiotic supplementation on intestinal microbiota and physiological conditions of Nile tilapia (Oreochromis niloticus) under waterborne cadmium exposure. Antonie van Leeuwenhoek 2017;110:501–513. 10.1007/s10482-016-0819-x

30. Xia J et al. Effects of short term lead exposure on gut microbiota and hepatic metabolism in adult zebrafish. Comparative Biochemistry and Physiology Part C: Toxicology & Pharmacology 2018;209:1–8. 10.1016/j.cbpc.2018.03.007

31. Luan Y et al. The Fish Microbiota: Research Progress and Potential Applications. Engineering 2023;29:137–146. 10.1016/j.eng.2022.12.011

32. Nayak SK. Role of gastrointestinal microbiota in fish. Aquaculture Research 2010;41:1553– 1573. 10.1111/j.1365-2109.2010.02546.x

33. Tran NT et al. Altered gut microbiota associated with intestinal disease in grass carp (Ctenopharyngodon idellus). World J Microbiol Biotechnol 2018;34:71. 10.1007/s11274-018-2447-2

34. Bozzi D et al. Salmon gut microbiota correlates with disease infection status: potential for monitoring health in farmed animals. Animal Microbiome 2021;3:30. 10.1186/s42523-021-00096-2

35. Li T et al. Alterations of the gut microbiome of largemouth bronze gudgeon (Coreius guichenoti) suffering from furunculosis. Sci Rep 2016;6:30606. 10.1038/srep30606

36. Aliko V et al. Antioxidant defense system, immune response and erythron profile modulation in gold fish, Carassius auratus, after acute manganese treatment. Fish Shellfish Immunol 2018;76:101–109. 10.1016/j.fsi.2018.02.042

37. Petitjean Q et al. Adaptive plastic responses to metal contamination in a multistress context: a field experiment in fish. Environ Sci Pollut Res 2023. 10.1007/s11356-023-26189-w

38. Claus SP, Guillou H, Ellero-Simatos S. The gut microbiota: a major player in the toxicity of environmental pollutants? NPJ Biofilms Microbiomes 2016;2:16003. 10.1038/npjbiofilms.2016.3

39. Evariste L et al. Gut microbiota of aquatic organisms: A key endpoint for ecotoxicological studies. Environ Pollut 2019;248:989–999. 10.1016/j.envpol.2019.02.101

40. Potts LD et al. Chronic Environmental Perturbation Influences Microbial Community Assembly Patterns. Environ Sci Technol 2022;56:2300–2311. 10.1021/acs.est.1c05106

41. Lavrinienko A et al. Applying the Anna Karenina principle for wild animal gut microbiota: Temporal stability of the bank vole gut microbiota in a disturbed environment. Journal of Animal Ecology 2020;89:2617–2630. 10.1111/1365-2656.13342

42. Zaneveld JR, McMinds R, Vega Thurber R. Stress and stability: applying the Anna Karenina principle to animal microbiomes. Nat Microbiol 2017;2:17121. 10.1038/nmicrobiol.2017.121

43. Rausch P et al. Expression of the Blood-Group-Related Gene B4galnt2 Alters Susceptibility to Salmonella Infection. PLOS Pathogens 2015;11:e1005008. 10.1371/journal.ppat.1005008

44. White J et al. Editorial: Impact of anthropogenic environmental changes on animal microbiomes. Frontiers in Ecology and Evolution 2023;11.

45. Williams K et al. Effects of subchronic exposure of silver nanoparticles on intestinal microbiota and gut-associated immune responses in the ileum of Sprague-Dawley rats. Nanotoxicology 2015;9:279–289. 10.3109/17435390.2014.921346

46. Zhang L et al. Persistent Organic Pollutants Modify Gut Microbiota-Host Metabolic Homeostasis in Mice Through Aryl Hydrocarbon Receptor Activation. Environ Health Perspect 2015;123:679–688. 10.1289/ehp.1409055

47. Xie L et al. Intestinal flora variation reflects the short-term damage of microplastic to the intestinal tract in mice. Ecotoxicology and Environmental Safety 2022;246:114194. 10.1016/j.ecoenv.2022.114194

48. Teffera M et al. Diverse mechanisms by which chemical pollutant exposure alters gut microbiota metabolism and inflammation. Environment International 2024;190:108805. 10.1016/j.envint.2024.108805

49. He X et al. Structural and functional alterations of gut microbiome in mice induced by chronic cadmium exposure. Chemosphere 2020;246:125747. 10.1016/j.chemosphere.2019.125747

50. Stott B, Elsdon JWV, Johnston JAA. Homing behaviour in gudgeon (Gobio gobio, (L.)). Animal Behaviour 1963;11:93–96. 10.1016/0003-3472(63)90015-0

51. Stott B. The Movements and Population Densities of Roach (Rutilus rutilus (L.)) and Gudgeon (Gobio gobio (L.)) in the River Mole. Journal of Animal Ecology 1967;36:407–423. 10.2307/2922

52. Petitjean Q. Variabilité de réponse aux stress multiples chez le goujon (Gobio occitaniae). 2019. These de doctorat, Toulouse 3.

53. Petitjean Q et al. Direct and indirect effects of multiple environmental stressors on fish health in human-altered rivers. Science of The Total Environment 2020;742:140657. 10.1016/j.scitotenv.2020.140657

54. Petitjean Q et al. Dose- and time-dependent effects of an immune challenge on fish across biological levels. Journal of Experimental Zoology Part A: Ecological and Integrative Physiology 2020;n/a. 10.1002/jez.2430

55. Boyer F et al. obitools: a unix-inspired software package for DNA metabarcoding. Mol Ecol Resour 2016;16:176–182. 10.1111/1755-0998.12428

56. Mercier C, et al. SUMATRA and SUMACLUST: fast and exact comparison and clustering of sequences. 2013. 2013.

57. Quast C et al. The SILVA ribosomal RNA gene database project: improved data processing and web-based tools. Nucleic Acids Research 2013;41:D590–D596. 10.1093/nar/gks1219

58. Bokulich NA et al. Quality-filtering vastly improves diversity estimates from Illumina amplicon sequencing. Nat Methods 2013;10:57–59. 10.1038/nmeth.2276

59. Zinger L et al. metabaR: An r package for the evaluation and improvement of DNA metabarcoding data quality. Methods in Ecology and Evolution 2021;12:586–592. 10.1111/2041-210X.13552

60. Nearing JT et al. Microbiome differential abundance methods produce different results across 38 datasets. Nat Commun 2022;13:342. 10.1038/s41467-022-28034-z

61. Cappellato M, Baruzzo G, Di Camillo B. Investigating differential abundance methods in microbiome data: A benchmark study. PLoS Comput Biol 2022;18:e1010467. 10.1371/journal.pcbi.1010467

62. McMurdie PJ, Holmes S. Waste Not, Want Not: Why Rarefying Microbiome Data Is Inadmissible. PLOS Computational Biology 2014;10:e1003531. 10.1371/journal.pcbi.1003531

63. Schloss PD. Waste not, want not: Revisiting the analysis that called into question the practice of rarefaction. Microbiology, 2023.

64. Gloor GB et al. Microbiome Datasets Are Compositional: And This Is Not Optional. Frontiers in Microbiology 2017;8.

65. Oksanen J et al. vegan: Community Ecology Package. 2019. 2019.

66. Kembel SW et al. Picante: R tools for integrating phylogenies and ecology. Bioinformatics 2010;26:1463–1464. 10.1093/bioinformatics/btq166

67. Wright ES. Using DECIPHER v2.0 to Analyze Big Biological Sequence Data in R. The R Journal 2016;8:352–359.

68. Schliep KP. phangorn: phylogenetic analysis in R. Bioinformatics 2011;27:592–593. 10.1093/bioinformatics/btq706

69. Bates D et al. Fitting Linear Mixed-Effects Models Using lme4. Journal of Statistical Software 2015;67(1):1–48. 10.18637/jss.v067.i01

70. Bartoń K. MuMIn: Multi-Model Inference. 2023. 2023.

71. McMurdie PJ, Holmes S. phyloseq: An R Package for Reproducible Interactive Analysis and Graphics of Microbiome Census Data. PLOS ONE 2013;8:e61217. 10.1371/journal.pone.0061217

72. Douglas GM et al. PICRUSt2 for prediction of metagenome functions. Nat Biotechnol 2020;38:685–688. 10.1038/s41587-020-0548-6

73. McDonald D et al. The Biological Observation Matrix (BIOM) format or: how I learned to stop worrying and love the ome-ome. GigaScience 2012;1:2047–217X-1–7. 10.1186/2047-217X-1-7

74. Caspi R et al. The MetaCyc database of metabolic pathways and enzymes and the BioCyc collection of pathway/genome databases. Nucleic Acids Res 2016;44:D471–480. 10.1093/nar/gkv1164

75. Zhou H et al. LinDA: linear models for differential abundance analysis of microbiome compositional data. Genome Biol 2022;23:1–23. 10.1186/s13059-022-02655-5

76. Benjamini Y, Hochberg Y. Controlling the False Discovery Rate: A Practical and Powerful Approach to Multiple Testing. Journal of the Royal Statistical Society Series B (Methodological) 1995;57:289–300. 10.1111/j.2517-6161.1995.tb02031.x

77. Brila I et al. Low-level environmental metal pollution is associated with altered gut microbiota of a wild rodent, the bank vole (*Myodes glareolus*). Science of The Total Environment 2021;790:148224. 10.1016/j.scitotenv.2021.148224

78. Kakade A et al. Long-term exposure of high concentration heavy metals induced toxicity, fatality, and gut microbial dysbiosis in common carp, *Cyprinus carpio*. Environmental Pollution 2020;266:115293. 10.1016/j.envpol.2020.115293

79. Matsuzaki R et al. Pesticide exposure and the microbiota-gut-brain axis. ISME J 2023;17:1153– 1166. 10.1038/s41396-023-01450-9

80. Rosenfeld CS. Gut Dysbiosis in Animals Due to Environmental Chemical Exposures. Front Cell Infect Microbiol 2017;7:396. 10.3389/fcimb.2017.00396

81. Jousset A et al. Where less may be more: how the rare biosphere pulls ecosystems strings. ISME J 2017;11:853–862. 10.1038/ismej.2016.174

82. von Hertzen L, Hanski I, Haahtela T. Natural immunity. Biodiversity loss and inflammatory diseases are two global megatrends that might be related. EMBO Rep 2011;12:1089–1093. 10.1038/embor.2011.195

83. Haahtela T. A biodiversity hypothesis. Allergy 2019;74:1445–1456. 10.1111/all.13763

84. Roslund MI et al. Biodiversity intervention enhances immune regulation and health-associated commensal microbiota among daycare children. Science Advances 2020;6:eaba2578. 10.1126/sciadv.aba2578

85. Kim PS et al. Host habitat is the major determinant of the gut microbiome of fish. Microbiome 2021;9:166. 10.1186/s40168-021-01113-x

86. Sullam KE et al. Environmental and ecological factors that shape the gut bacterial communities of fish: a meta-analysis. Mol Ecol 2012;21:3363–3378. 10.1111/j.1365-294X.2012.05552.x

87. Maritan E et al. The role of animal hosts in shaping gut microbiome variation. Philosophical Transactions of the Royal Society B: Biological Sciences 2024;379:20230071. 10.1098/rstb.2023.0071

88. Pepke ML, Hansen SB, Limborg MT. Unraveling host regulation of gut microbiota through the epigenome–microbiome axis. Trends in Microbiology 2024. 10.1016/j.tim.2024.05.006

89. Parthasarathy A et al. A Three-Ring Circus: Metabolism of the Three Proteogenic Aromatic Amino Acids and Their Role in the Health of Plants and Animals. Front Mol Biosci 2018;5. 10.3389/fmolb.2018.00029

90. Aquili L. The Role of Tryptophan and Tyrosine in Executive Function and Reward Processing. Int J Tryptophan Res 2020;13:1178646920964825. 10.1177/1178646920964825

91. Khan N, Deschaux P. Role of serotonin in fish immunomodulation. J Exp Biol 1997;200:1833– 1838. 10.1242/jeb.200.13.1833

92. de Abreu MS et al. Dopamine and serotonin mediate the impact of stress on cleaner fish cooperative behavior. Horm Behav 2020;125:104813. 10.1016/j.yhbeh.2020.104813

93. Leal E et al. Effects of dopaminergic system activation on feeding behavior and growth performance of the sea bass (Dicentrarchus labrax): a self-feeding approach. Horm Behav 2013;64:113–121. 10.1016/j.yhbeh.2013.05.008

94. Schultz W. Getting Formal with Dopamine and Reward. Neuron 2002;36:241–263. 10.1016/S0896-6273(02)00967-4

95. Friedman ES et al. Microbes vs. chemistry in the origin of the anaerobic gut lumen. Proceedings of the National Academy of Sciences 2018;115:4170–4175. 10.1073/pnas.1718635115

96. Jurburg SD et al. Synthesis of recovery patterns in microbial communities across environments. Microbiome 2024;12:79. 10.1186/s40168-024-01802-3

97. Alberdi A, Martin Bideguren G, Aizpurua O. Diversity and compositional changes in the gut microbiota of wild and captive vertebrates: a meta-analysis. Sci Rep 2021;11:22660. 10.1038/s41598-021-02015-6

98. Pimentel T et al. Bacterial communities 16S rDNA fingerprinting as a potential tracing tool for cultured seabass Dicentrarchus labrax. Sci Rep 2017;7:11862. 10.1038/s41598-017-11552-y

99. Côte J et al. Changes in fish skin microbiota along gradients of eutrophication in human-altered rivers. FEMS Microbiol Ecol 2022;98:fiac006. 10.1093/femsec/fiac006

100. Mullens N et al. Anna Karenina as a promoter of microbial diversity in the cosmopolitan agricultural pest Zeugodacus cucurbitae (Diptera, Tephritidae). PLoS ONE 2024;19:e0300875. 10.1371/journal.pone.0300875

