## Supplementary Materials for "Experimental metal contamination reduces gut microbiota diversity and alters its composition and function in wild-caught fish"

Table Of Content

### Supplementary material 1. List of the six artefactual MOTUs identified. Contamination mostly occurred during DNA extraction.

| Phylum | Class | Order | Family | Genus | Species |
| --- | --- | --- | --- | --- | --- |
| Actinobacteriota | Actinobacteria | Actinomycetales | [Actinomycetaceae](mailto:Actinomycetaceae@family) | [Actinomyces](mailto:Actinomyces@genus%22;1) | NA |
| Firmicutes | [Bacilli](mailto:Bacilli@class) | [Lactobacillales](mailto:Lactobacillales@order) | Carnobacteriaceae | Trichococcus | uncultured bacterium |
| Firmicutes | Bacilli | Lactobacillales | Lactobacillaceae | Paucilactobacillus | uncultured Lactobacillus sp. |
| Proteobacteria | Alphaproteobacteria | Acetobacterales | Acetobacteraceae | Acidiphilium | uncultured bacterium |
| Proteobacteria | Alphaproteobacteria | Rhodobacterales | Rhodobacteraceae | Paracoccus | uncultured bacterium |
| Proteobacteria | Gammaproteobacteria | Burkholderiales | Neisseriaceae | Uruburuella | uncultured bacterium |

### Supplementary material 2. Data Normalization

As current literature suggests that the choice of normalization method can greatly impact the interpretation of 16S DNA sequence read counts [e.g., [3–7]], we performed α- and β-diversity analyses on the original dataset as well as on a set of 5 commonly used normalization methods (i.e., proportion, cumulative sum scaling-CSS, rarefaction, deseq, trimmed mean of M value-TMM) and their log-transformed (log2(x+1)) version to verify the consistency of our results. Here, we report the Chisq or F value (depending on the test conducted) for treatments and/or covariates retained in the best final models (selected using AICc) for each transformation. Missing treatment or covariate for a given transformation means it was excluded from the best final model. The stars indicate a significant effect of the treatment or covariate on either α-diversity (Supplementary Material 2A), β-diversity (Supplementary Material 2B), or betaDisper (Supplementary Material 2C). A brief discussion of the differences among normalization methods and indices is provided in Supplementary Material 2D.

#### Supplementary material 2A. α-diversity

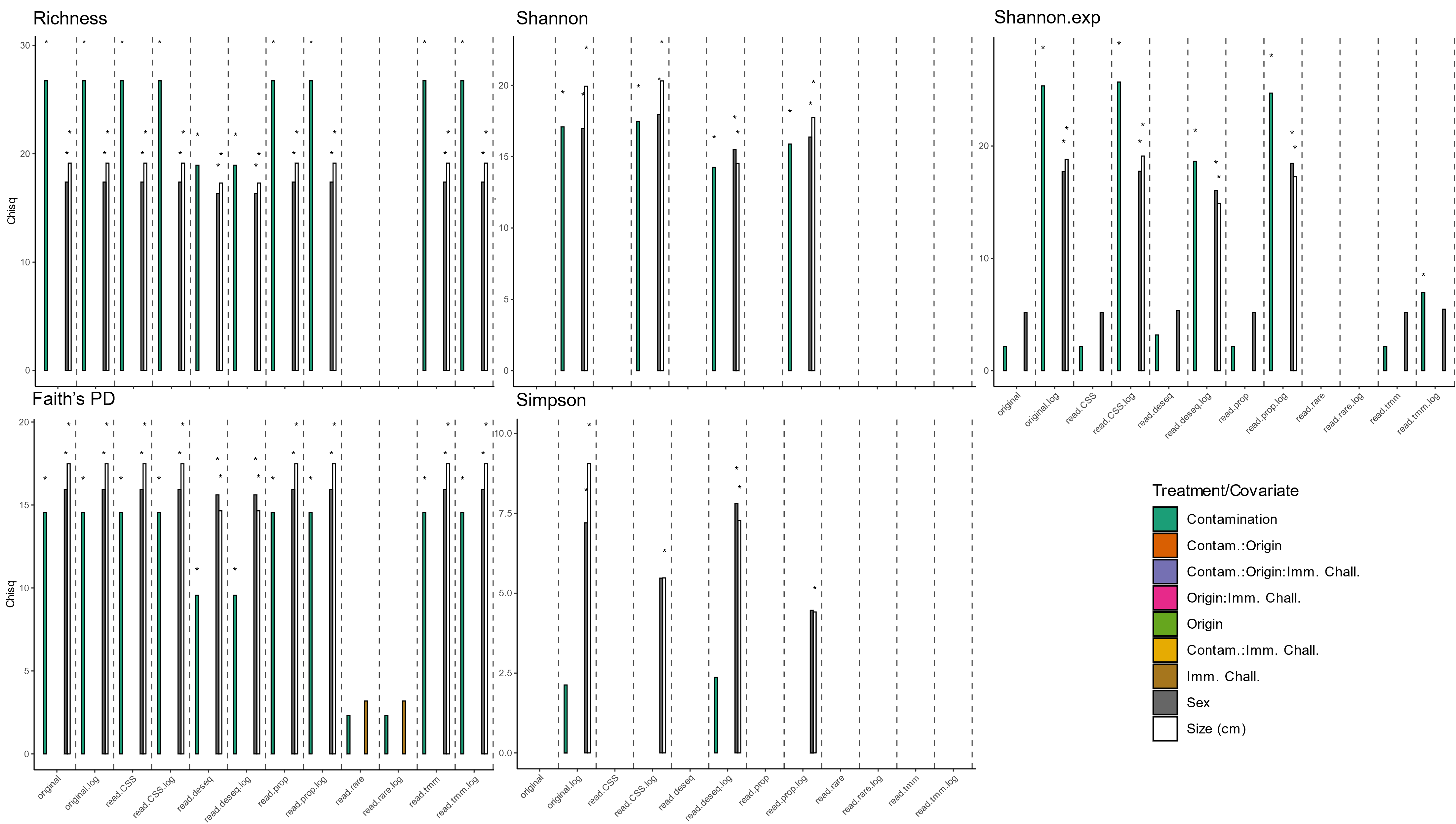
For α-diversity indices (i.e., Richness, Shannon, Shannon.exp, Faith’s PD, and Simpson), we can see that for most transformations (except rarefying) and considered indices (except Simpson), the effect of the contamination treatment and the sex and size of the fish remain in the final models and are mostly significant, indicating that, in our study, the results are consistent across normalization methods (see plots below).

#### Supplementary material 2B. β-diversity

For β-diversity distances (i.e., PCOA coordinates from the first and second axes for Bray-Curtis, Jaccard, Robust Aitchison, Hellinger, and Weighed and unweighed Unifrac), we can see that for most transformations (except rarefying and for all normalization with Unifrac distance for PCOA axis 2), the effect of the contamination treatment and the sex and size of the fish remain in the final models and are mostly significant, indicating that, in our study, the results are consistent across normalization methods (see plots below).

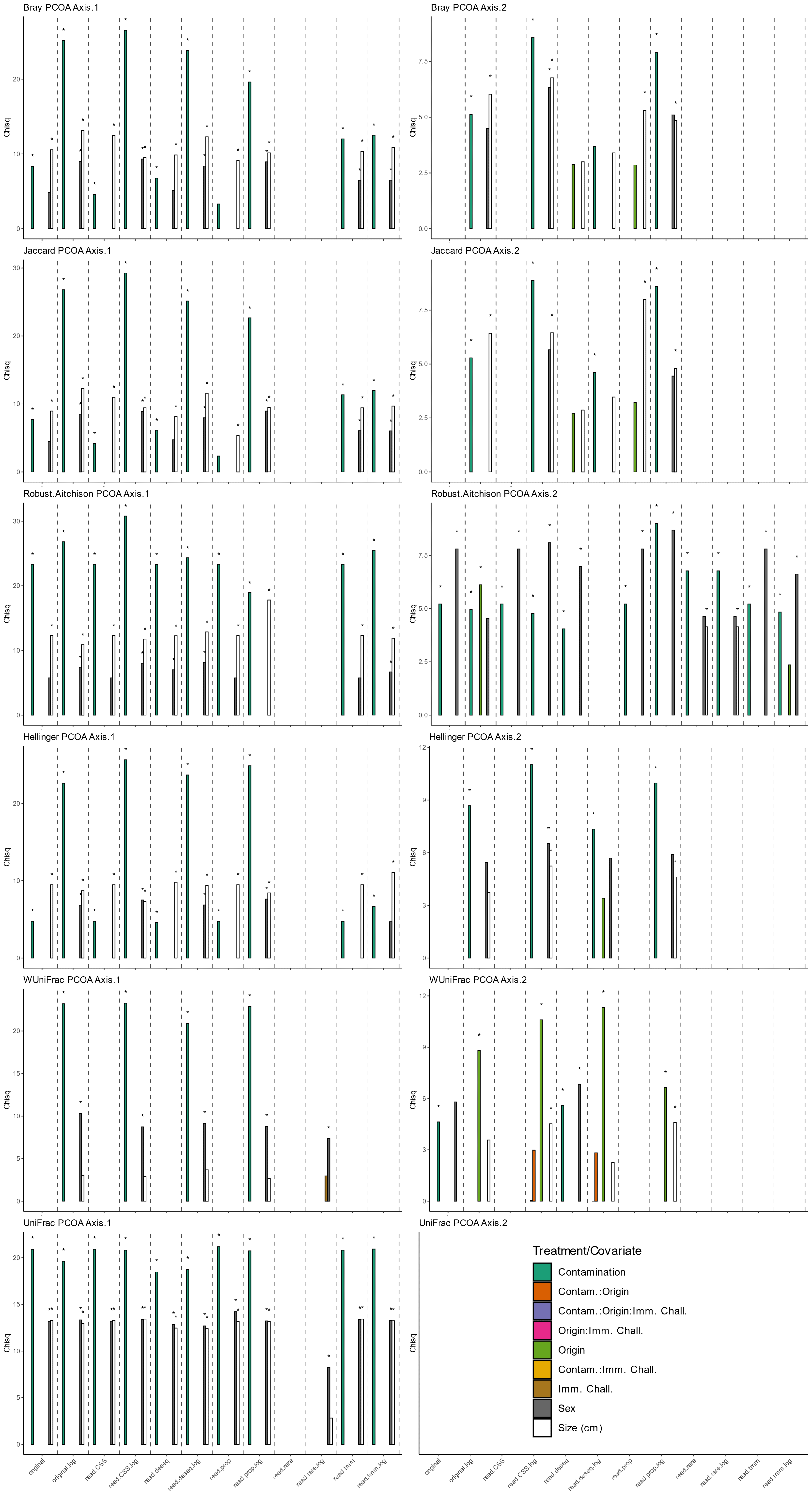

#### Supplementary material 2C. Betadisper

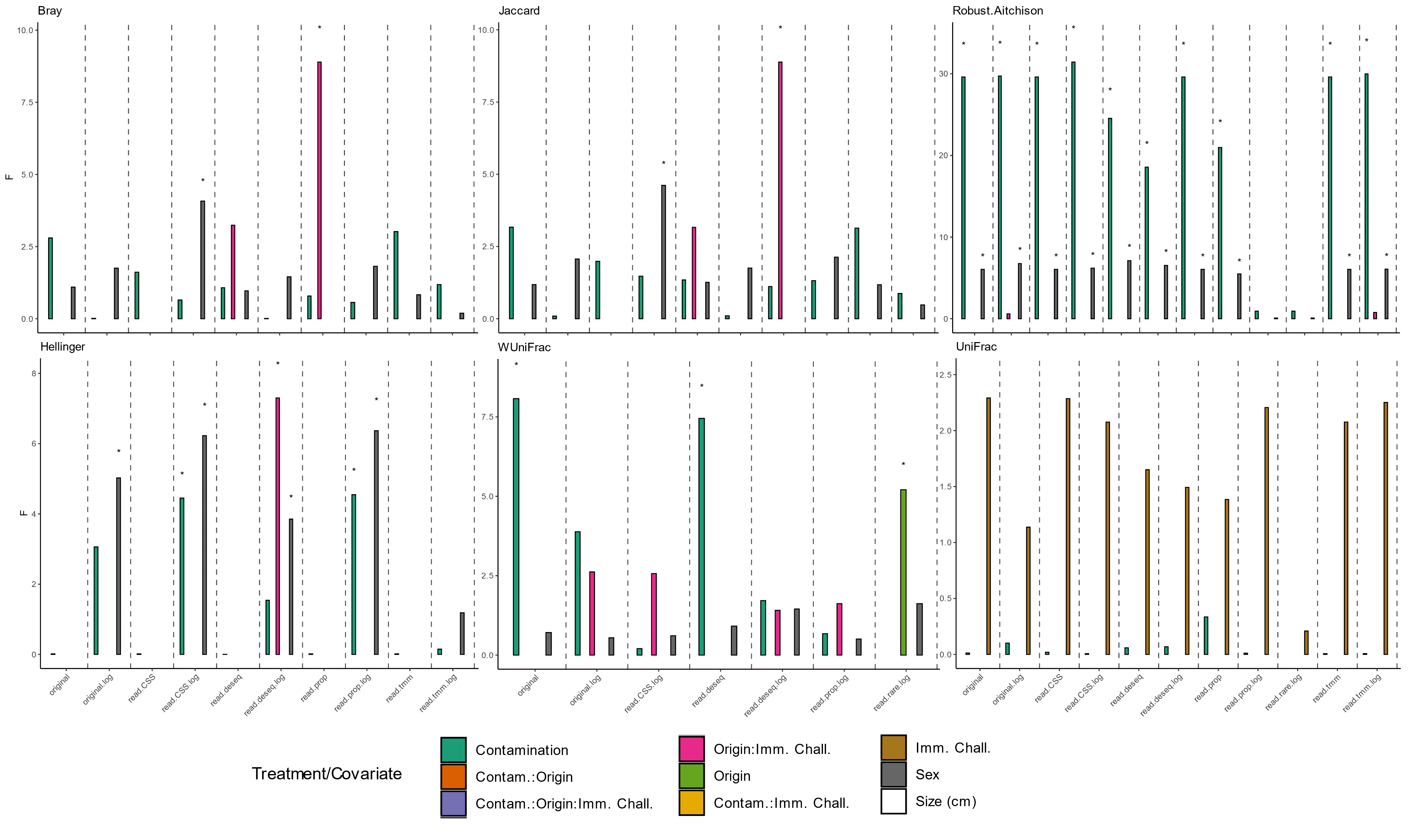
For betadisper performed on dissimilarity matrices computed according to several distances (i.e., Bray-Curtis, Jaccard, Robust Aitchison, Hellinger, and Weighed and unweighed Unifrac), we can see that for most transformations, the effect of the treatment is not significant although some treatment and covariate remain in the best final model, except for Robust Aitchison distance where contamination treatment and sex of the fish are significant, indicating that, in our study, the results are moslty consistent across normalization methods (see plots below).

#### Supplementary material 2D. Discrepancies among normalization methods and dissimilarity indices

Overall, our study's use of multiple normalization methods and computation of various dissimilarity (β-diversity) indices highlight the critical role of choosing appropriate methods for tackling questions investigating shifts in microbial communities.

Despite our findings suggest robust consistency across most normalization methods for α- and β-diversity analyses, we observed notable exceptions where the choice of method or index may lead to different interpretations of the data. For example, rarefaction generally showed less consistency in capturing the effects of treatments on diversity indices compared to other methods, such as log-transformed original data, CSS, or DESeq, which aligns with discussions in recent studies questioning the reliability of rarefaction [7].

Specifically, while most methods yielded consistent results in β-diversity indices, the Robust Aitchison and Weighted UniFrac indices sometimes produced different outcomes compared to other indices. For instance, Robust Aitchison and Weighted UniFrac allowed the detection of significant dissimilarity regarding taxonomic composition on PCOA axis 2 (see Supplementary Material 5), contrary to other dissimilarity indices such as Bray or Jaccard, although models explained a low amount of variance (R2m Robust.Aitchison = 0.038; R2m WUniFrac = 0.028, see supplementary material 5). Hence, these indices indicated minor inconsistencies when detecting changes due to treatments. While these discrepancies could be due to different mathematical formulations and assumptions (e.g., standard vs. compositional approach, see [6]), they also highlight the need to compare the results among various indices before concluding.

Interestingly, when examining the dispersion (i.e., variance) of the community composition among treatments (betadisper analyses), we noticed that group dispersion was mostly unaffected across indices. However, exceptions were observed, for instance, with the Robust Aitchison index for contamination treatment and sex, suggesting that the compositional approach [6], through Aitchison distance, might be more sensitive to specific types of community shifts.

To conclude, the few discrepancies observed among normalization methods and indices underscore the importance of selecting appropriate normalization and dissimilarity measures tailored to microbiome research's specific questions and data characteristics. Although our results were mostly consistent, the highlighted inconsistencies reveal that such methodological choices can influence the detection and interpretation of potential alteration of the microbial communities. While relying on every existing normalization and indices when analyzing microbiome data is not the best way to reach robust results and conclusions, the selection of the approaches should be done with caution to ensure the reliability of the results. Hence, without appropriate knowledge, checking for consistency across methods and indices could help provide a clearer view of what is happening in the dataset. Accordingly, future studies might benefit from carefully evaluating these methods to ensure accurate ecological inferences, supporting a broader call for standardized reporting and method selection in microbiome research.

### Supplementary material 3. Phylogenetic tree construction

The phylogenetic tree was constructed using the phangorn R package [1]. First, DNA sequences were aligned using the DECIPHER R package [2]; we built the initial tree based on neighbor-joining tree estimation and Hamming distance. Second, we tested for all nucleotide substitution models included in the phangorn package and retrieved the best one (transition model – TIM3+G(4)+I) according to the likelihood of the phylogenetic tree given sequence alignment and tested models. Parameters of the selected model were then optimized for topology, base frequencies, gamma rate, proportion of variable size, rate matrix, and edge lengths. We finally tested whether the optimized tree was better than the raw one using Shimodaira-Hasegawa Test for phylogenetic inferences. The best phylogenetic tree is displayed below. Here, tree tips correspond to family names, and colors correspond to phylum.

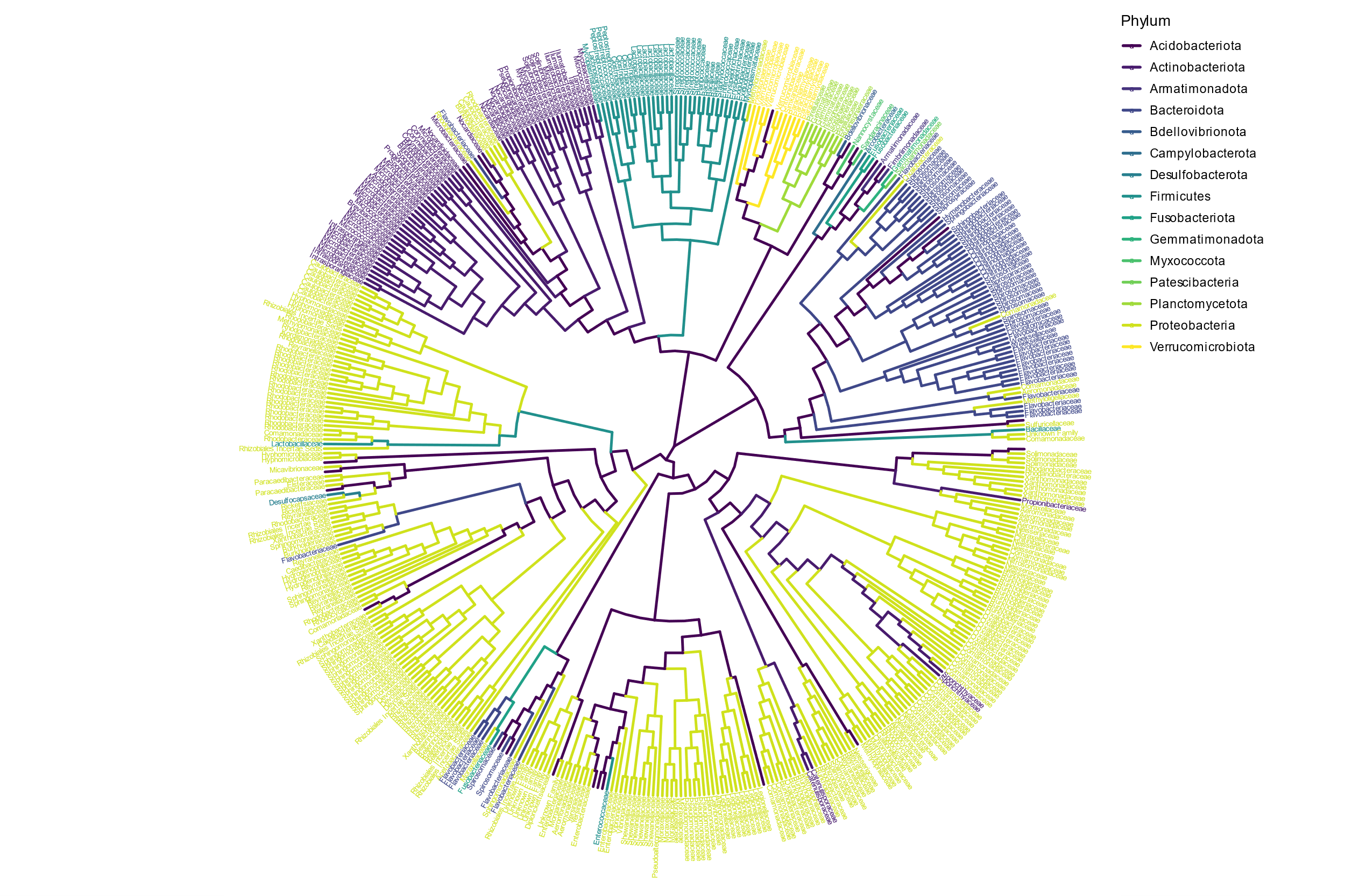

### Supplementary material 4. Summary of the results of the linear models (LM) testing the effects of metal contamination (NC vs. C), immune challenge (PBS vs. AMIX), and the fish origin (NC vs. C) on α-diversity indices (Richness, Shannon, exp(Shannon), Simpson and Faith's PD) in the water performed on original log-transformed data. Sample size and multiple and adjusted R square are reported as n, Mult. R2, and Adj. R2, respectively.

Here, contamination by trace metal elements significantly reduces the α-diversity of bacteria in water, regardless of the α-diversity index.

|  | Estimate | Std. Error | t value | df | SSq | F_value | p.value |
| --- | --- | --- | --- | --- | --- | --- | --- |
| ***Richness \| n = 32 \| Mult. R2 = 0.302 \| Adj. R2 = 0.134*** | | | | | | | |
| Intercept | 256 | 18.5 | 13.8 | 1 | 252000 | 190 | <0.0001 |
| Contamination (NC) | 37.5 | 13.1 | 2.85 | 1 | 10800 | 8.13 | <0.01 |
| PopAUSCOR | -3.36 | 20.4 | -0.165 | 4 | 3170 | 0.597 | 0.668 |
| PopCELCAB | -0.984 | 24.5 | -0.0402 | 4 | 3170 | 0.597 | 0.668 |
| PopFARM | -10.9 | 21.4 | -0.512 | 4 | 3170 | 0.597 | 0.668 |
| PopRIOU | 17.9 | 21.3 | 0.837 | 4 | 3170 | 0.597 | 0.668 |
| Imm. Chall. (PBS) | 0.334 | 13.4 | 0.0249 | 1 | 0.823 | 0.00062 | 0.98 |
| Residuals | NA | NA | NA | 25 | 33200 | NA | NA |
| ***Shannon \| n = 32 \| Mult. R2 = 0.314 \| Adj. R2 = 0.149*** | | | | | | | |
| Intercept | 5.29 | 0.0783 | 67.6 | 1 | 108 | 4570 | <0.0001 |
| Contamination (NC) | 0.159 | 0.0555 | 2.86 | 1 | 0.194 | 8.18 | <0.01 |
| PopAUSCOR | -0.0308 | 0.0863 | -0.356 | 4 | 0.0675 | 0.713 | 0.591 |
| PopCELCAB | -0.0388 | 0.104 | -0.375 | 4 | 0.0675 | 0.713 | 0.591 |
| PopFARM | -0.0766 | 0.0903 | -0.848 | 4 | 0.0675 | 0.713 | 0.591 |
| PopRIOU | 0.0565 | 0.0901 | 0.627 | 4 | 0.0675 | 0.713 | 0.591 |
| Imm. Chall. (PBS) | 0.00212 | 0.0566 | 0.0374 | 1 | 0.0000331 | 0.0014 | 0.97 |
| Residuals | NA | NA | NA | 25 | 0.592 | NA | NA |
| ***Shannon.exp \| n = 32 \| Mult. R2 = 0.33 \| Adj. R2 = 0.169*** | | | | | | | |
| Intercept | 199 | 15.6 | 12.7 | 1 | 152000 | 161 | <0.0001 |
| Contamination (NC) | 33.5 | 11.1 | 3.02 | 1 | 8610 | 9.12 | <0.01 |
| PopAUSCOR | -4.42 | 17.2 | -0.256 | 4 | 2580 | 0.682 | 0.611 |
| PopCELCAB | -3.51 | 20.7 | -0.17 | 4 | 2580 | 0.682 | 0.611 |
| PopFARM | -13 | 18 | -0.723 | 4 | 2580 | 0.682 | 0.611 |
| PopRIOU | 13.4 | 18 | 0.742 | 4 | 2580 | 0.682 | 0.611 |
| Imm. Chall. (PBS) | 1.2 | 11.3 | 0.106 | 1 | 10.6 | 0.0112 | 0.917 |
| Residuals | NA | NA | NA | 25 | 23600 | NA | NA |
| ***Simpson \| n = 32 \| Mult. R2 = 0.277 \| Adj. R2 = 0.103*** | | | | | | | |
| Intercept | 0.994 | 0.000563 | 1770 | 1 | 3.81 | 3120000 | <0.0001 |
| Contamination (NC) | 0.000965 | 0.000399 | 2.42 | 1 | 0.00000715 | 5.84 | <0.05 |
| PopAUSCOR | -0.000271 | 0.000621 | -0.437 | 4 | 0.00000414 | 0.846 | 0.509 |
| PopCELCAB | -0.00036 | 0.000745 | -0.483 | 4 | 0.00000414 | 0.846 | 0.509 |
| PopFARM | -0.000667 | 0.000649 | -1.03 | 4 | 0.00000414 | 0.846 | 0.509 |
| PopRIOU | 0.000372 | 0.000648 | 0.574 | 4 | 0.00000414 | 0.846 | 0.509 |
| Imm. Chall. (PBS) | 0.0000116 | 0.000407 | 0.0286 | 1 | 0.000000001 | 0.000819 | 0.977 |
| Residuals | NA | NA | NA | 25 | 0.0000306 | NA | NA |
| ***PD \| n = 32 \| Mult. R2 = 0.33 \| Adj. R2 = 0.169*** | | | | | | | |
| Intercept | 33.1 | 1.61 | 20.5 | 1 | 4230 | 421 | <0.0001 |
| Contamination (NC) | 3.79 | 1.14 | 3.31 | 1 | 110 | 11 | <0.01 |
| PopAUSCOR | 0.11 | 1.78 | 0.0619 | 4 | 10.9 | 0.271 | 0.894 |
| PopCELCAB | -0.328 | 2.13 | -0.154 | 4 | 10.9 | 0.271 | 0.894 |
| PopFARM | 0.248 | 1.86 | 0.133 | 4 | 10.9 | 0.271 | 0.894 |
| PopRIOU | 1.41 | 1.86 | 0.76 | 4 | 10.9 | 0.271 | 0.894 |
| Imm. Chall. (PBS) | -0.271 | 1.17 | -0.232 | 1 | 0.543 | 0.054 | 0.818 |
| Residuals | NA | NA | NA | 25 | 251 | NA | NA |

### Supplementary material 5. Summary of the results of the best (selected using AICc) final linear mixed models (LMM) testing the effects of metal contamination (NC vs. C), immune challenge (PBS vs. AMIX), fish origin (NC vs. C) and covariates (fish size and sex) on PCOA scores (second axis) representing β-diversity indices (Bray Curtis, Jaccard, Robust Aitchison, Hellinger, Weighed Unifrac and Unweighted Unifrac) on **fish gut microbiota community** (left) and **inferred functional composition** (from picrust2, right) performed on original log-transformed data. Sample size and marginal and conditional R square are reported as n, R2m, and R2c, respectively. Unifrac and weighed Unifrac distances were not computed for functional composition as they rely on phylogenetic distances among taxa.

|  | Taxonomic composition | | | | | | Functional composition | | | | | |
| --- | --- | --- | --- | --- | --- | --- | --- | --- | --- | --- | --- | --- |
|  | Estimate | Std. Error | t value | df | Chisq | p.value | Estimate | Std. Error | t value | df | Chisq | p.value |
| ***Bray \| n = 307 \| R2m = 0.0395 \| R2c = 0.485*** | | | | | | | ***Bray \| n = 307 \| R2m = 0.0182 \| R2c = 0.0707*** | | | | | |
| Intercept | -0.0831 | 0.0718 | -1.16 | 1 | 1.34 | 0.247 | -0.000765 | 0.00591 | -0.129 | 1 | 0.0168 | 0.897 |
| Contamination (NC) | -0.0496 | 0.0219 | -2.26 | 1 | 5.12 | <0.05 | -0.00913 | 0.0058 | -1.57 | 1 | 2.48 | 0.116 |
| Origin (NC) | NA | NA | NA | NA | NA | NA | 0.00967 | 0.00627 | 1.54 | 1 | 2.38 | 0.123 |
| Sex (I) | 0.0104 | 0.0277 | 0.374 | 2 | 4.49 | 0.106 |  |  |  |  |  |  |
| Sex (M) | -0.0371 | 0.0194 | -1.91 | 2 | 4.49 | 0.106 |  |  |  |  |  |  |
| Size (cm) | 0.0144 | 0.00585 | 2.46 | 1 | 6.03 | <0.05 |  |  |  |  |  |  |
| ***Jaccard \| n = 307 \| R2m = 0.0348 \| R2c = 0.476*** | | | | | | | ***Jaccard \| n = 307 \| R2m = 0.0482 \| R2c = 0.221*** | | | | | |
| Intercept | -0.0899 | 0.0636 | -1.41 | 1 | 1.99 | 0.158 | -0.0413 | 0.0301 | -1.37 | 1 | 1.89 | 0.17 |
| Contamination (NC) | -0.0495 | 0.0216 | -2.3 | 1 | 5.28 | <0.05 | NA | NA | NA | NA | NA | NA |
| Origin (NC) | NA | NA | NA | NA | NA | NA | -0.0349 | 0.0177 | -1.97 | 1 | 3.88 | <0.05 |
| Size (cm) | 0.0136 | 0.00536 | 2.53 | 1 | 6.42 | <0.05 | 0.00621 | 0.00282 | 2.2 | 1 | 4.86 | <0.05 |
| ***Robust.Aitchison \| n = 307 \| R2m = 0.0383 \| R2c = 0.669*** | | | | | | | ***Robust.Aitchison \| n = 307 \| R2m = 0 \| R2c = 0.252*** | | | | | |
| Intercept | -0.184 | 0.309 | -0.595 | 1 | 0.354 | 0.552 | -0.147 | 0.283 | -0.519 | 1 | 0.269 | 0.604 |
| Contamination (NC) | 0.254 | 0.114 | 2.23 | 1 | 4.96 | <0.05 |  |  |  |  |  |  |
| Origin (NC) | -0.318 | 0.128 | -2.47 | 1 | 6.12 | <0.05 |  |  |  |  |  |  |
| Sex (I) | -0.000497 | 0.129 | -0.00384 | 2 | 4.54 | 0.103 |  |  |  |  |  |  |
| Sex (M) | 0.191 | 0.0929 | 2.06 | 2 | 4.54 | 0.103 |  |  |  |  |  |  |
| ***Hellinger \| n = 307 \| R2m = 0.0589 \| R2c = 0.344*** | | | | | | | ***Hellinger \| n = 307 \| R2m = 0 \| R2c = 0.564*** | | | | | |
| Intercept | -0.0758 | 0.0906 | -0.837 | 1 | 0.701 | 0.402 | -0.00773 | 0.0172 | -0.45 | 1 | 0.202 | 0.653 |
| Contamination (NC) | -0.0894 | 0.0304 | -2.95 | 1 | 8.68 | <0.01 |  |  |  |  |  |  |
| Sex (I) | 0.00337 | 0.039 | 0.0864 | 2 | 5.44 | 0.0658 |  |  |  |  |  |  |
| Sex (M) | -0.0607 | 0.0273 | -2.22 | 2 | 5.44 | 0.0658 |  |  |  |  |  |  |
| Size (cm) | 0.0158 | 0.00822 | 1.93 | 1 | 3.72 | 0.0539 |  |  |  |  |  |  |
| ***WUniFrac \| n = 307 \| R2m = 0.0284 \| R2c = 0.464*** | | | | | | |  |  |  |  |  |  |
| Intercept | -0.0341 | 0.0242 | -1.41 | 1 | 1.98 | 0.159 |  |  |  |  |  |  |
| Origin (NC) | -0.0235 | 0.00791 | -2.97 | 1 | 8.82 | <0.01 |  |  |  |  |  |  |
| Size (cm) | 0.0038 | 0.00201 | 1.89 | 1 | 3.57 | 0.0587 |  |  |  |  |  |  |
| ***UniFrac \| n = 307 \| R2m = 0 \| R2c = 0.706*** | | | | | | |  |  |  |  |  |  |
| Intercept | -0.0248 | 0.0467 | -0.53 | 1 | 0.281 | 0.596 |  |  |  |  |  |  |

### Supplementary material 6. Summary of the results of the linear models (LM) testing the effects of metal contamination (NC vs. C), immune challenge (PBS vs. AMIX), fish site of origin (FARM, CELCAB, ARIMAS, AUSCOR, RIOU) on PCOA scores (first axis: left, second axis: right) representing β-diversity indices (Bray Curtis, Jaccard, Robust Aitchison, Hellinger, Weighed Unifrac and Unweighted Unifrac) for **water samples** performed on original log-transformed data. Sample size and multiple and adjusted R square are reported as n, Mult. R2, and Adj. R2, respectively.

|  | PCOA Axis 1 | | | | | | | PCOA Axis 2 | | | | | | |
| --- | --- | --- | --- | --- | --- | --- | --- | --- | --- | --- | --- | --- | --- | --- |
|  | Estimate | Std. Error | t value | df | SSq | F_value | p.value | Estimate | Std. Error | t value | df | SSq | F_value | p.value |
| ***Bray \| n = 32 \| Mult. R2 = 0.954 \| Adj. R2 = 0.943*** | | | | | | | | ***Bray \| n = 32 \| Mult. R2 = 0.309 \| Adj. R2 = 0.143*** | | | | | | |
| Intercept | -0.164 | 0.019 | -8.6 | 1 | 0.103 | 74 | <0.0001 | -0.0753 | 0.0564 | -1.33 | 1 | 0.0219 | 1.78 | 0.194 |
| Contamination (NC) | 0.301 | 0.0135 | 22.3 | 1 | 0.696 | 499 | <0.0001 | -0.00324 | 0.04 | -0.081 | 1 | 0.0000807 | 0.00657 | 0.936 |
| PopAUSCOR | -0.027 | 0.021 | -1.29 | 4 | 0.00647 | 1.16 | 0.353 | -0.00222 | 0.0622 | -0.0356 | 4 | 0.109 | 2.21 | 0.0969 |
| PopCELCAB | 0.0128 | 0.0252 | 0.507 | 4 | 0.00647 | 1.16 | 0.353 | 0.156 | 0.0746 | 2.09 | 4 | 0.109 | 2.21 | 0.0969 |
| PopFARM | -0.00018 | 0.0219 | -0.00818 | 4 | 0.00647 | 1.16 | 0.353 | 0.0343 | 0.0651 | 0.528 | 4 | 0.109 | 2.21 | 0.0969 |
| PopRIOU | -0.0201 | 0.0219 | -0.917 | 4 | 0.00647 | 1.16 | 0.353 | 0.11 | 0.0649 | 1.69 | 4 | 0.109 | 2.21 | 0.0969 |
| Imm. Chall. (PBS) | 0.0103 | 0.0137 | 0.747 | 1 | 0.00078 | 0.559 | 0.462 | 0.061 | 0.0408 | 1.5 | 1 | 0.0275 | 2.24 | 0.147 |
| Residuals | NA | NA | NA | 25 | 0.0349 | NA | NA | NA | NA | NA | 25 | 0.307 | NA | NA |
| ***Jaccard \| n = 32 \| Mult. R2 = 0.958 \| Adj. R2 = 0.948*** | | | | | | | | ***Jaccard \| n = 32 \| Mult. R2 = 0.34 \| Adj. R2 = 0.182*** | | | | | | |
| Intercept | -0.206 | 0.0222 | -9.28 | 1 | 0.165 | 86.2 | <0.0001 | -0.0986 | 0.068 | -1.45 | 1 | 0.0375 | 2.1 | 0.159 |
| Contamination (NC) | 0.369 | 0.0158 | 23.4 | 1 | 1.04 | 546 | <0.0001 | -0.0191 | 0.0482 | -0.397 | 1 | 0.00281 | 0.158 | 0.695 |
| PopAUSCOR | -0.0326 | 0.0245 | -1.33 | 4 | 0.0164 | 2.15 | 0.104 | 0.0038 | 0.0749 | 0.0507 | 4 | 0.177 | 2.48 | 0.0695 |
| PopCELCAB | 0.0345 | 0.0294 | 1.17 | 4 | 0.0164 | 2.15 | 0.104 | 0.211 | 0.0899 | 2.34 | 4 | 0.177 | 2.48 | 0.0695 |
| PopFARM | 0.0111 | 0.0257 | 0.433 | 4 | 0.0164 | 2.15 | 0.104 | 0.0667 | 0.0784 | 0.851 | 4 | 0.177 | 2.48 | 0.0695 |
| PopRIOU | -0.0205 | 0.0256 | -0.799 | 4 | 0.0164 | 2.15 | 0.104 | 0.14 | 0.0782 | 1.79 | 4 | 0.177 | 2.48 | 0.0695 |
| Imm. Chall. (PBS) | 0.0136 | 0.0161 | 0.845 | 1 | 0.00136 | 0.714 | 0.406 | 0.0838 | 0.0491 | 1.71 | 1 | 0.0519 | 2.91 | 0.1 |
| Residuals | NA | NA | NA | 25 | 0.0478 | NA | NA | NA | NA | NA | 25 | 0.446 | NA | NA |
| ***Robust.Aitchison \| n = 32 \| Mult. R2 = 0.806 \| Adj. R2 = 0.76*** | | | | | | | | ***Robust.Aitchison \| n = 32 \| Mult. R2 = 0.464 \| Adj. R2 = 0.336*** | | | | | | |
| Intercept | -4.77 | 0.875 | -5.45 | 1 | 87.7 | 29.7 | <0.0001 | -0.624 | 1.24 | -0.502 | 1 | 1.5 | 0.252 | 0.62 |
| Contamination (NC) | 5.51 | 0.621 | 8.88 | 1 | 233 | 78.9 | <0.0001 | -3.02 | 0.881 | -3.43 | 1 | 69.9 | 11.7 | <0.01 |
| PopAUSCOR | -0.0432 | 0.965 | -0.0448 | 4 | 55.1 | 4.66 | <0.01 | 0.417 | 1.37 | 0.305 | 4 | 41.6 | 1.75 | 0.171 |
| PopCELCAB | 3.99 | 1.16 | 3.45 | 4 | 55.1 | 4.66 | <0.01 | 3.19 | 1.64 | 1.94 | 4 | 41.6 | 1.75 | 0.171 |
| PopFARM | 1.86 | 1.01 | 1.84 | 4 | 55.1 | 4.66 | <0.01 | 1.92 | 1.43 | 1.34 | 4 | 41.6 | 1.75 | 0.171 |
| PopRIOU | 1.27 | 1.01 | 1.26 | 4 | 55.1 | 4.66 | <0.01 | 2.52 | 1.43 | 1.76 | 4 | 41.6 | 1.75 | 0.171 |
| Imm. Chall. (PBS) | 1.12 | 0.632 | 1.78 | 1 | 9.35 | 3.16 | 0.0875 | 1.91 | 0.897 | 2.13 | 1 | 26.9 | 4.52 | <0.05 |
| Residuals | NA | NA | NA | 25 | 73.9 | NA | NA | NA | NA | NA | 25 | 149 | NA | NA |
| ***Hellinger \| n = 32 \| Mult. R2 = 0.957 \| Adj. R2 = 0.947*** | | | | | | | | ***Hellinger \| n = 32 \| Mult. R2 = 0.469 \| Adj. R2 = 0.341*** | | | | | | |
| Intercept | -0.266 | 0.0296 | -9 | 1 | 0.273 | 80.9 | <0.0001 | -0.11 | 0.0698 | -1.57 | 1 | 0.0465 | 2.47 | 0.129 |
| Contamination (NC) | 0.487 | 0.021 | 23.2 | 1 | 1.82 | 538 | <0.0001 | -0.0335 | 0.0495 | -0.675 | 1 | 0.00859 | 0.456 | 0.506 |
| PopAUSCOR | -0.0404 | 0.0326 | -1.24 | 4 | 0.0113 | 0.839 | 0.514 | -0.0251 | 0.077 | -0.326 | 4 | 0.319 | 4.24 | <0.01 |
| PopCELCAB | 0.016 | 0.0391 | 0.409 | 4 | 0.0113 | 0.839 | 0.514 | 0.286 | 0.0924 | 3.1 | 4 | 0.319 | 4.24 | <0.01 |
| PopFARM | -0.00554 | 0.0341 | -0.162 | 4 | 0.0113 | 0.839 | 0.514 | 0.0942 | 0.0806 | 1.17 | 4 | 0.319 | 4.24 | <0.01 |
| PopRIOU | -0.018 | 0.034 | -0.529 | 4 | 0.0113 | 0.839 | 0.514 | 0.13 | 0.0804 | 1.62 | 4 | 0.319 | 4.24 | <0.01 |
| Imm. Chall. (PBS) | 0.0154 | 0.0214 | 0.72 | 1 | 0.00175 | 0.519 | 0.478 | 0.116 | 0.0505 | 2.3 | 1 | 0.0999 | 5.3 | <0.05 |
| Residuals | NA | NA | NA | 25 | 0.0845 | NA | NA | NA | NA | NA | 25 | 0.471 | NA | NA |
| ***WUniFrac \| n = 32 \| Mult. R2 = 0.846 \| Adj. R2 = 0.809*** | | | | | | | | ***WUniFrac \| n = 32 \| Mult. R2 = 0.403 \| Adj. R2 = 0.26*** | | | | | | |
| Intercept | -0.0571 | 0.0121 | -4.72 | 1 | 0.0126 | 22.3 | <0.0001 | 0.0174 | 0.0143 | 1.22 | 1 | 0.00117 | 1.49 | 0.234 |
| Contamination (NC) | 0.096 | 0.00857 | 11.2 | 1 | 0.0707 | 125 | <0.0001 | 0.0128 | 0.0101 | 1.26 | 1 | 0.00125 | 1.58 | 0.22 |
| PopAUSCOR | -0.0121 | 0.0133 | -0.91 | 4 | 0.00115 | 0.509 | 0.73 | 0.00334 | 0.0158 | 0.212 | 4 | 0.0083 | 2.63 | 0.0584 |
| PopCELCAB | 0.00199 | 0.016 | 0.124 | 4 | 0.00115 | 0.509 | 0.73 | -0.0456 | 0.0189 | -2.41 | 4 | 0.0083 | 2.63 | 0.0584 |
| PopFARM | 0.00254 | 0.0139 | 0.182 | 4 | 0.00115 | 0.509 | 0.73 | -0.0235 | 0.0165 | -1.43 | 4 | 0.0083 | 2.63 | 0.0584 |
| PopRIOU | 0.000309 | 0.0139 | 0.0222 | 4 | 0.00115 | 0.509 | 0.73 | -0.0162 | 0.0165 | -0.984 | 4 | 0.0083 | 2.63 | 0.0584 |
| Imm. Chall. (PBS) | 0.0129 | 0.00873 | 1.47 | 1 | 0.00122 | 2.17 | 0.153 | -0.0255 | 0.0103 | -2.47 | 1 | 0.00481 | 6.1 | <0.05 |
| Residuals | NA | NA | NA | 25 | 0.0141 | NA | NA | NA | NA | NA | 25 | 0.0197 | NA | NA |
| ***UniFrac \| n = 32 \| Mult. R2 = 0.909 \| Adj. R2 = 0.888*** | | | | | | | | ***UniFrac \| n = 32 \| Mult. R2 = 0.112 \| Adj. R2 = -0.101*** | | | | | | |
| Intercept | -0.121 | 0.0196 | -6.18 | 1 | 0.0569 | 38.2 | <0.0001 | -0.00189 | 0.0386 | -0.0489 | 1 | 0.0000138 | 0.00239 | 0.961 |
| Contamination (NC) | 0.214 | 0.0139 | 15.4 | 1 | 0.352 | 236 | <0.0001 | 0.0067 | 0.0274 | 0.245 | 1 | 0.000345 | 0.0598 | 0.809 |
| PopAUSCOR | -0.0111 | 0.0217 | -0.515 | 4 | 0.00191 | 0.32 | 0.862 | -0.00624 | 0.0426 | -0.146 | 4 | 0.0179 | 0.776 | 0.551 |
| PopCELCAB | 0.00291 | 0.026 | 0.112 | 4 | 0.00191 | 0.32 | 0.862 | 0.0266 | 0.0511 | 0.521 | 4 | 0.0179 | 0.776 | 0.551 |
| PopFARM | -0.0175 | 0.0227 | -0.773 | 4 | 0.00191 | 0.32 | 0.862 | -0.0367 | 0.0446 | -0.823 | 4 | 0.0179 | 0.776 | 0.551 |
| PopRIOU | 0.000117 | 0.0226 | 0.00518 | 4 | 0.00191 | 0.32 | 0.862 | 0.028 | 0.0445 | 0.63 | 4 | 0.0179 | 0.776 | 0.551 |
| Imm. Chall. (PBS) | 0.0173 | 0.0142 | 1.22 | 1 | 0.00221 | 1.48 | 0.235 | -0.00357 | 0.0279 | -0.128 | 1 | 0.0000944 | 0.0164 | 0.899 |
| Residuals | NA | NA | NA | 25 | 0.0372 | NA | NA | NA | NA | NA | 25 | 0.144 | NA | NA |

### Supplementary material 7. Results of the Envfit approach testing the effects of metal contamination (NC vs. C), immune challenge (PBS vs. AMIX), fish origin (NC vs. C), and covariates (fish size and sex) on the dissimilarity of fish gut microbiota community (left table) and water microbiota community (right table) using original log-transformed data.

Here, the treatments (Contamination and immune challenge) and the level of contamination at the populations’ origin site affect the dispersion of individuals in the PCOA space (i.e., ordination scores), whatever the index considered (except for the Unifrac index, where fish origin is marginally significant) in fish gut samples. On the contrary, only the contamination treatment affects the dispersion of individuals in the PCOA space in water samples. In addition, in fish gut the covariates (fish sex and size) also affect the dispersion of individuals in the PCOA space (except for the weighted Unifrac index, where fish size is not significant). However, the correlation coefficient is relatively low, whatever the variable and the index considered. It suggests that while treatment affects the dissimilarity of fish gut microbiota community, they might not be the primary driver of community dissimilarity contrary to what is observed in the water where correlation coefficients are high (above 40%).

| ***Fish gut (MOTUs)*** | | |
| --- | --- | --- |
|  | r2 | p.value |
| ***Bray*** | | |
| Contamination | 0.05070 | <0.01 |
| Origin | 0.02360 | <0.01 |
| Imm. Chall. | 0.03610 | <0.01 |
| Sex | 0.02250 | <0.05 |
| Size (cm) | 0.06190 | <0.01 |
| ***Jaccard*** | | |
| Contamination | 0.05210 | <0.01 |
| Origin | 0.02310 | <0.01 |
| Imm. Chall. | 0.03550 | <0.01 |
| Sex | 0.02140 | 0.01 |
| Size (cm) | 0.06120 | <0.01 |
| ***Robust.Aitchison*** | | |
| Contamination | 0.06540 | <0.01 |
| Origin | 0.02120 | <0.01 |
| Imm. Chall. | 0.02420 | <0.01 |
| Sex | 0.02210 | <0.05 |
| Size (cm) | 0.03350 | <0.01 |
| ***Hellinger*** | | |
| Contamination | 0.05060 | <0.01 |
| Origin | 0.02720 | <0.01 |
| Imm. Chall. | 0.04060 | <0.01 |
| Sex | 0.01890 | <0.05 |
| Size (cm) | 0.04770 | <0.01 |
| ***WUniFrac*** | | |
| Contamination | 0.04740 | <0.01 |
| Origin | 0.02200 | <0.01 |
| Imm. Chall. | 0.02940 | <0.01 |
| Sex | 0.02040 | <0.05 |
| Size (cm) | 0.00391 | 0.532 |
| ***UniFrac*** | | |
| Contamination | 0.04060 | <0.01 |
| Origin | 0.00937 | 0.057 |
| Imm. Chall. | 0.04420 | <0.01 |
| Sex | 0.02410 | <0.01 |
| Size (cm) | 0.04460 | <0.01 |

| ***Fish gut (Functions)*** | | |
| --- | --- | --- |
|  | r2 | p.value |
| ***Bray*** | | |
| Contamination | 0.05540 | <0.01 |
| Origin | 0.01810 | <0.01 |
| Imm. Chall. | 0.01340 | <0.05 |
| Sex | 0.04090 | <0.01 |
| Size (cm) | 0.02980 | <0.05 |
| ***Jaccard*** | | |
| Contamination | 0.05680 | <0.01 |
| Origin | 0.01690 | <0.05 |
| Imm. Chall. | 0.01340 | <0.05 |
| Sex | 0.03680 | <0.01 |
| Size (cm) | 0.03010 | <0.05 |
| ***Robust.Aitchison*** | | |
| Contamination | 0.04810 | <0.01 |
| Origin | 0.03470 | <0.01 |
| Imm. Chall. | 0.02520 | <0.01 |
| Sex | 0.02030 | <0.05 |
| Size (cm) | 0.01470 | 0.108 |
| ***Hellinger*** | | |
| Contamination | 0.04250 | <0.01 |
| Origin | 0.01060 | <0.05 |
| Imm. Chall. | 0.00816 | 0.075 |
| Sex | 0.02870 | <0.01 |
| Size (cm) | 0.12800 | <0.01 |

| ***Water samples (MOTUs)*** | | |
| --- | --- | --- |
|  | r2 | p.value |
| ***Bray*** | | |
| Contamination | 0.5970 | <0.01 |
| Imm. Chall. | 0.0398 | 0.281 |
| Pop | 0.0959 | 0.678 |
| ***Jaccard*** | | |
| Contamination | 0.5930 | <0.01 |
| Imm. Chall. | 0.0429 | 0.262 |
| Pop | 0.1050 | 0.603 |
| ***Robust.Aitchison*** | | |
| Contamination | 0.4660 | <0.01 |
| Imm. Chall. | 0.0595 | 0.15 |
| Pop | 0.1330 | 0.416 |
| ***Hellinger*** | | |
| Contamination | 0.6580 | <0.01 |
| Imm. Chall. | 0.0474 | 0.23 |
| Pop | 0.1140 | 0.511 |
| ***WUniFrac*** | | |
| Contamination | 0.6120 | <0.01 |
| Imm. Chall. | 0.0673 | 0.135 |
| Pop | 0.0716 | 0.8 |
| ***UniFrac*** | | |
| Contamination | 0.6440 | <0.01 |
| Imm. Chall. | 0.0328 | 0.359 |
| Pop | 0.0355 | 0.973 |

### Supplementary material 8. Results of the betadisper approach followed by permutation test (999 permutations) testing the effects of metal contamination (NC vs. C), immune challenge (PBS vs. AMIX), fish origin (NC vs. C), and covariates (fish size and sex) on the dispersion homogeneity in fish gut microbiota community using original log-transformed data.

***Taxonomic composition***

|  | Df | Sum Sq | Mean Sq | F | N.Perm | Pr(>F) |
| --- | --- | --- | --- | --- | --- | --- |
| ***Bray*** | | | | | | |
| Contamination | 1 | 2.75e-05 | 2.75e-05 | 0.0142 | 999 | 0.895 |
| Residuals | 305 | 5.92e-01 | 1.94e-03 |  |  |  |
| Sex | 2 | 5.87e-03 | 2.93e-03 | 1.75 | 999 | 0.19 |
| Residuals | 304 | 5.08e-01 | 1.67e-03 |  |  |  |
| ***Jaccard*** | | | | | | |
| Contamination | 1 | 1.78e-04 | 1.78e-04 | 0.0915 | 999 | 0.765 |
| Residuals | 305 | 5.95e-01 | 1.95e-03 |  |  |  |
| Sex | 2 | 6.24e-03 | 3.12e-03 | 2.07 | 999 | 0.116 |
| Residuals | 304 | 4.58e-01 | 1.51e-03 |  |  |  |
| ***Robust.Aitchison*** | | | | | | |
| Contamination | 1 | 5.98e+01 | 5.98e+01 | 29.7 | 999 | <0.01 |
| Residuals | 305 | 6.14e+02 | 2.01e+00 |  |  |  |
| Sex | 2 | 3.07e+01 | 1.54e+01 | 6.76 | 999 | <0.01 |
| Residuals | 304 | 6.91e+02 | 2.27e+00 |  |  |  |
| Origin | 1 | 1.44e+00 | 1.44e+00 | 0.601 | 999 | 0.458 |
| Residuals | 305 | 7.31e+02 | 2.40e+00 |  |  |  |
| ***Hellinger*** | | | | | | |
| Contamination | 1 | 1.79e-02 | 1.79e-02 | 3.06 | 999 | 0.091 |
| Residuals | 305 | 1.79e+00 | 5.86e-03 |  |  |  |
| Sex | 2 | 4.16e-02 | 2.08e-02 | 5.03 | 999 | <0.01 |
| Residuals | 304 | 1.26e+00 | 4.14e-03 |  |  |  |
| ***WUniFrac*** | | | | | | |
| Contamination | 1 | 1.65e-03 | 1.65e-03 | 3.88 | 999 | <0.05 |
| Residuals | 305 | 1.30e-01 | 4.25e-04 |  |  |  |
| Sex | 2 | 4.76e-04 | 2.38e-04 | 0.545 | 999 | 0.6 |
| Residuals | 304 | 1.33e-01 | 4.37e-04 |  |  |  |
| Origin | 1 | 1.10e-03 | 1.10e-03 | 2.62 | 999 | 0.116 |
| Residuals | 305 | 1.28e-01 | 4.20e-04 |  |  |  |
| ***UniFrac*** | | | | | | |
| Contamination | 1 | 1.72e-04 | 1.72e-04 | 0.102 | 999 | 0.754 |
| Residuals | 305 | 5.16e-01 | 1.69e-03 |  |  |  |
| Sex | 2 | 3.57e-03 | 1.78e-03 | 1.14 | 999 | 0.283 |
| Residuals | 304 | 4.77e-01 | 1.57e-03 |  |  |  |

| ***Functional composition*** | | | | | | |
| --- | --- | --- | --- | --- | --- | --- |
|  | Df | Sum Sq | Mean Sq | F | N.Perm | Pr(>F) |
| ***Bray*** | | | | | | |
| Contamination | 1 | 4.34e-03 | 0.004340 | 2.2 | 999 | 0.147 |
| Residuals | 305 | 6.02e-01 | 0.001970 |  |  |  |
| Sex | 2 | 1.85e-03 | 0.000923 | 0.483 | 999 | 0.619 |
| Residuals | 304 | 5.81e-01 | 0.001910 |  |  |  |
| Origin | 1 | 9.25e-03 | 0.009250 | 4.88 | 999 | <0.05 |
| Residuals | 305 | 5.78e-01 | 0.001890 |  |  |  |
| ***Jaccard*** | | | | | | |
| Contamination | 1 | 9.81e-03 | 0.009810 | 2.44 | 999 | 0.142 |
| Residuals | 305 | 1.23e+00 | 0.004030 |  |  |  |
| Sex | 2 | 1.76e-03 | 0.000881 | 0.224 | 999 | 0.805 |
| Residuals | 304 | 1.20e+00 | 0.003930 |  |  |  |
| Origin | 1 | 2.38e-02 | 0.023800 | 6.19 | 999 | <0.05 |
| Residuals | 305 | 1.17e+00 | 0.003840 |  |  |  |
| ***Robust.Aitchison*** | | | | | | |
| Contamination | 1 | 3.06e-01 | 0.306000 | 0.263 | 999 | 0.625 |
| Residuals | 305 | 3.55e+02 | 1.160000 |  |  |  |
| Sex | 2 | 6.87e-01 | 0.344000 | 0.306 | 999 | 0.727 |
| Residuals | 304 | 3.41e+02 | 1.120000 |  |  |  |
| Origin | 1 | 6.26e-01 | 0.626000 | 0.579 | 999 | 0.439 |
| Residuals | 305 | 3.30e+02 | 1.080000 |  |  |  |
| ***Hellinger*** | | | | | | |
| Contamination | 1 | 3.99e-03 | 0.003990 | 1.34 | 999 | 0.261 |
| Residuals | 305 | 9.07e-01 | 0.002970 |  |  |  |
| Sex | 2 | 2.04e-02 | 0.010200 | 3.71 | 999 | <0.05 |
| Residuals | 304 | 8.38e-01 | 0.002760 |  |  |  |

### Supplementary material 9 Summary of the results of the linear models for differential abundance analysis (LINDA) testing the effects of metal contamination (NC vs. C), immune challenge (PBS vs. AMIX), and the fish origin (NC vs. C) on fish gut microbiome composition at the family level. Analysis was performed on original log-transformed data. Only the taxa that have been significantly (adjusted p < 0.05 and |log2FoldChange| > 0.5) differentially abundant among treatments are displayed. Taxa for which relative abundance changes according to contamination treatment in both water and gut are highlighted in bold.

| ***Differential abundance of Taxa at the family level - LINDA results for fish gut*** | | | | | | | |
| --- | --- | --- | --- | --- | --- | --- | --- |
|  | baseMean | log2FoldChange | lfcSE | stat | df | pvalue | padj |
| ***Contamination (NC vs. C)*** | | | | | | | |
| **Aeromonadaceae** | **5100** | **-2.30** | **0.520** | **-4.5** | **6** | **3.4e-05** | **2.5e-04** |
| Beijerinckiaceae | 770 | -1.80 | 0.190 | -9.4 | 30 | 0.0e+00 | 0.0e+00 |
| Chitinibacteraceae | 1100 | -1.20 | 0.320 | -3.9 | 6 | 2.4e-04 | 1.3e-03 |
| Chromobacteriaceae | 240 | -0.76 | 0.160 | -4.7 | 6 | 1.4e-05 | 1.2e-04 |
| Comamonadaceae | 4500 | -0.77 | 0.230 | -3.4 | 6 | 1.2e-03 | 5.1e-03 |
| Devosiaceae | 320 | -0.75 | 0.190 | -4.0 | 5 | 2.1e-04 | 1.2e-03 |
| Enterococcaceae | 1600 | -0.60 | 0.200 | -3.0 | 5 | 4.5e-03 | 1.7e-02 |
| Iamiaceae | 950 | -2.30 | 0.350 | -6.6 | 6 | 0.0e+00 | 1.0e-07 |
| **Ilumatobacteraceae** | **300** | **-1.10** | **0.140** | **-8.3** | **6** | **0.0e+00** | **0.0e+00** |
| Legionellaceae | 730 | -1.20 | 0.280 | -4.3 | 6 | 5.7e-05 | 3.9e-04 |
| **Microbacteriaceae** | **3500** | **-0.80** | **0.160** | **-4.9** | **6** | **5.4e-06** | **4.9e-05** |
| **Microtrichaceae** | **260** | **-0.80** | **0.130** | **-6.2** | **6** | **0.0e+00** | **5.0e-07** |
| **Mycobacteriaceae** | **670** | **-0.88** | **0.230** | **-3.8** | **4** | **4.0e-04** | **1.9e-03** |
| **Neisseriaceae** | **320** | **-0.65** | **0.180** | **-3.7** | **30** | **3.1e-04** | **1.5e-03** |
| Nocardioidaceae | 340 | -0.53 | 0.180 | -3.0 | 6 | 4.2e-03 | 1.6e-02 |
| Pirellulaceae | 380 | -1.00 | 0.160 | -6.6 | 7 | 0.0e+00 | 1.0e-07 |
| **Pseudomonadaceae** | **1500** | **-1.00** | **0.240** | **-4.1** | **5** | **1.2e-04** | **7.5e-04** |
| **Rhizobiaceae** | **1100** | **-3.20** | **0.290** | **-11.0** | **5** | **0.0e+00** | **0.0e+00** |
| **Rhizobiales Incertae Sedis** | **320** | **-1.30** | **0.200** | **-6.3** | **5** | **0.0e+00** | **5.0e-07** |
| Rhodobacteraceae | 4200 | -1.30 | 0.360 | -3.5 | 6 | 7.8e-04 | 3.3e-03 |
| **Solirubrobacteraceae** | **300** | **-1.10** | **0.120** | **-8.7** | **6** | **0.0e+00** | **0.0e+00** |
| **Unknown Family** | **350** | **-1.40** | **0.260** | **-5.6** | **5** | **8.0e-07** | **7.7e-06** |
| ***Imm. Chall. (PBS vs. AMIX)*** | | | | | | | |
| Pseudomonadaceae | 1500 | 1.20 | 0.270 | 4.6 | 6 | 2.2e-05 | 2.0e-03 |
| ***Sex (F vs. M)*** | | | | | | | |
| Beijerinckiaceae | 770 | 0.77 | 0.210 | 3.6 | 29 | 3.9e-04 | 5.8e-03 |
| Iamiaceae | 950 | 0.93 | 0.290 | 3.2 | 29 | 1.7e-03 | 2.2e-02 |
| Microbacteriaceae | 3500 | 0.64 | 0.180 | 3.6 | 29 | 3.5e-04 | 5.8e-03 |
| Rhizobiaceae | 1100 | 1.30 | 0.280 | 4.6 | 30 | 5.1e-06 | 4.6e-04 |
| Rhizobiales Incertae Sedis | 320 | 0.82 | 0.210 | 3.9 | 29 | 1.1e-04 | 3.8e-03 |
| Unknown Family | 350 | 0.69 | 0.180 | 3.8 | 28 | 2.0e-04 | 4.5e-03 |
| Xanthomonadaceae | 320 | 0.65 | 0.170 | 3.9 | 29 | 1.3e-04 | 3.8e-03 |
| ***Size (cm)*** | | | | | | | |
| Burkholderiaceae | 36000 | -0.52 | 0.120 | -4.3 | 25 | 2.3e-05 | 4.2e-04 |
| Enterococcaceae | 1600 | -0.66 | 0.098 | -6.7 | 28 | 0.0e+00 | 0.0e+00 |
| Moraxellaceae | 9000 | -0.91 | 0.260 | -3.5 | 29 | 4.7e-04 | 3.5e-03 |
| Nocardiaceae | 1600 | -0.69 | 0.160 | -4.4 | 24 | 2.0e-05 | 4.2e-04 |
| Pseudomonadaceae | 1500 | -0.51 | 0.120 | -4.1 | 24 | 5.6e-05 | 8.3e-04 |
| Reyranellaceae | 1000 | -0.68 | 0.190 | -3.6 | 23 | 3.6e-04 | 3.0e-03 |
| Rhodobacteraceae | 4200 | -0.65 | 0.190 | -3.5 | 28 | 5.8e-04 | 4.0e-03 |

### Supplementary material 10. Summary of the results of the linear models for differential abundance analysis (LINDA) testing the effects of metal contamination (NC vs. C), immune challenge (PBS vs. AMIX), and the fish origin (NC vs. C) on fish gut microbiome composition at the phylum level. Analysis was performed on original log-transformed data. Only the taxa that have been significantly (adjusted p < 0.05 and |log2FoldChange| > 0.5) differentially abundant among treatments are displayed. Taxa for which relative abundance changes according to contamination treatment in both water and gut are highlighted in bold.

| ***Differential abundance of Taxa at the phylum level - LINDA results for fish gut*** | | | | | | | |
| --- | --- | --- | --- | --- | --- | --- | --- |
|  | baseMean | log2FoldChange | lfcSE | stat | df | pvalue | padj |
| ***Contamination (NC vs. C)*** | | | | | | | |
| **Actinobacteriota** | **7700** | **-0.85** | **0.17** | **-4.9** | **6** | **8.1e-06** | **6.2e-05** |
| Planctomycetota | 190 | -1.00 | 0.21 | -4.8 | 6 | 9.5e-06 | 6.2e-05 |
| **Proteobacteria** | **85000** | **-0.78** | **0.20** | **-3.9** | **6** | **2.4e-04** | **7.6e-04** |
| ***Sex (F vs. M)*** | | | | | | | |
| Actinobacteriota | 7700 | 0.65 | 0.17 | 3.7 | 30 | 2.2e-04 | 1.5e-03 |
| Planctomycetota | 190 | 0.51 | 0.20 | 2.5 | 28 | 1.2e-02 | 3.0e-02 |
| Proteobacteria | 85000 | 0.71 | 0.20 | 3.5 | 30 | 5.6e-04 | 2.4e-03 |
| Verrucomicrobiota | 160 | 0.79 | 0.19 | 4.2 | 30 | 3.6e-05 | 4.6e-04 |
| ***Size (cm)*** | | | | | | | |
| Bacteroidota | 1200 | -0.52 | 0.14 | -3.8 | 27 | 1.6e-04 | 3.5e-04 |
| Proteobacteria | 85000 | -0.65 | 0.10 | -6.3 | 29 | 0.0e+00 | 0.0e+00 |

### Supplementary material 11. Summary of the results of the best (Selected using AICc) final linear mixed models (LMM) testing the effects of metal contamination (NC vs. C), immune challenge (PBS vs. AMIX), fish origin (NC vs. C) and covariates (fish size and sex) on Firmicutes/Bacteroidetes (F/B; left table) and the Proteobacteria/Bacteroidetes (P/B; right table) ratio.

| ***F/B ratio*** | | | | | | |
| --- | --- | --- | --- | --- | --- | --- |
|  | Estimate | Std. Error | t value | df | Chisq | p.value |
| ***Original \| n = 248 \| R2m = 0 \| R2c = 0.0259*** | | | | | | |
| Intercept | 65.6 | 43 | 1.53 | 1 | 2.33 | 0.127 |
| ***Original (log-transformed) \| n = 248 \| R2m = 0 \| R2c = 0.315*** | | | | | | |
| Intercept | 2.11 | 0.336 | 6.29 | 1 | 39.6 | <0.0001 |
| ***Proportion \| n = 248 \| R2m = 0 \| R2c = 0.0259*** | | | | | | |
| Intercept | 65.6 | 43 | 1.53 | 1 | 2.33 | 0.127 |
| ***Proportion (log-transformed) \| n = 248 \| R2m = 0 \| R2c = 0.124*** | | | | | | |
| Intercept | 1.53 | 0.158 | 9.71 | 1 | 94.2 | <0.0001 |
| ***CSS \| n = 248 \| R2m = 0 \| R2c = 0.0259*** | | | | | | |
| Intercept | 65.6 | 43 | 1.53 | 1 | 2.33 | 0.127 |
| ***CSS (log-transformed) \| n = 248 \| R2m = 0 \| R2c = 0.0382*** | | | | | | |
| Intercept | 1.4 | 0.141 | 9.95 | 1 | 99.1 | <0.0001 |
| ***DESEQ \| n = 248 \| R2m = 0 \| R2c = 0.0335*** | | | | | | |
| Intercept | 51.4 | 31 | 1.66 | 1 | 2.74 | 0.0979 |
| ***DESEQ (log-transformed) \| n = 248 \| R2m = 0 \| R2c = 0.0642*** | | | | | | |
| Intercept | 2.23 | 0.43 | 5.19 | 1 | 26.9 | <0.0001 |
| ***TMM \| n = 248 \| R2m = 0 \| R2c = 0.0259*** | | | | | | |
| Intercept | 65.6 | 43 | 1.53 | 1 | 2.33 | 0.127 |
| ***TMM (log-transformed) \| n = 248 \| R2m = 0 \| R2c = 0.0368*** | | | | | | |
| Intercept | 43.2 | 27.1 | 1.59 | 1 | 2.54 | 0.111 |

| ***P/B ratio*** | | | | | | |
| --- | --- | --- | --- | --- | --- | --- |
|  | Estimate | Std. Error | t value | df | Chisq | p.value |
| ***Original \| n = 306 \| R2m = 0 \| R2c = 0.191*** | | | | | | |
| Intercept | 0.0453 | 0.0116 | 3.9 | 1 | 15.2 | <0.0001 |
| ***Original (log-transformed) \| n = 306 \| R2m = 0 \| R2c = 0.379*** | | | | | | |
| Intercept | 0.349 | 0.0494 | 7.06 | 1 | 49.9 | <0.0001 |
| ***Proportion \| n = 306 \| R2m = 0 \| R2c = 0.191*** | | | | | | |
| Intercept | 0.0453 | 0.0116 | 3.9 | 1 | 15.2 | <0.0001 |
| ***Proportion (log-transformed) \| n = 306 \| R2m = 0 \| R2c = 0.27*** | | | | | | |
| Intercept | 0.459 | 0.0463 | 9.91 | 1 | 98.2 | <0.0001 |
| ***CSS \| n = 306 \| R2m = 0 \| R2c = 0.191*** | | | | | | |
| Intercept | 0.0453 | 0.0116 | 3.9 | 1 | 15.2 | <0.0001 |
| ***CSS (log-transformed) \| n = 306 \| R2m = 0 \| R2c = 0.279*** | | | | | | |
| Intercept | 0.512 | 0.0505 | 10.1 | 1 | 103 | <0.0001 |
| ***DESEQ \| n = 306 \| R2m = 0.0205 \| R2c = 0.192*** | | | | | | |
| Intercept | 0.0447 | 0.0156 | 2.86 | 1 | 8.17 | <0.01 |
| Contamination (NC) | -0.0226 | 0.0152 | -1.49 | 1 | 2.22 | 0.137 |
| Imm. Chall. (PBS) | 0.0225 | 0.016 | 1.41 | 1 | 2 | 0.158 |
| ***DESEQ (log-transformed) \| n = 306 \| R2m = 0 \| R2c = 0.359*** | | | | | | |
| Intercept | 0.352 | 0.045 | 7.83 | 1 | 61.2 | <0.0001 |
| ***TMM \| n = 306 \| R2m = 0 \| R2c = 0.191*** | | | | | | |
| Intercept | 0.0453 | 0.0116 | 3.9 | 1 | 15.2 | <0.0001 |
| ***TMM (log-transformed) \| n = 306 \| R2m = 0.0222 \| R2c = 0.203*** | | | | | | |
| Intercept | 0.0524 | 0.0171 | 3.07 | 1 | 9.44 | <0.01 |
| Contamination (NC) | -0.0252 | 0.0161 | -1.57 | 1 | 2.46 | 0.117 |
| Imm. Chall. (PBS) | 0.0246 | 0.017 | 1.45 | 1 | 2.1 | 0.147 |

### Supplementary material 12 Summary of the results of the linear models for differential abundance analysis (LINDA) testing the effects of metal contamination (NC vs. C), immune challenge (PBS vs. AMIX), and the fish origin (NC vs. C) on water (within fish tanks) microbiome composition at the family level. Analysis was performed on original log-transformed data. Only the taxa that have been significantly (adjusted p < 0.05 and |log2FoldChange| > 0.5) differentially abundant among treatments are displayed.Taxa for which relative abundance changes according to contamination treatment in both water and gut are highlighted in bold.

| ***Differential abundance of Taxa at the family level - LINDA results for water*** | | | | | | | |
| --- | --- | --- | --- | --- | --- | --- | --- |
|  | baseMean | log2FoldChange | lfcSE | stat | df | pvalue | padj |
| ***Contamination (NC vs. C)*** | | | | | | | |
| Acetobacteraceae | 5 | 1.30 | 0.41 | 3.1 | 2 | 4.1e-03 | 1.4e-02 |
| Acidothermaceae | 120 | -3.20 | 0.52 | -6.1 | 2 | 1.4e-06 | 1.4e-05 |
| **Aeromonadaceae** | **110** | **-2.60** | **0.90** | **-2.9** | **2** | **7.5e-03** | **2.1e-02** |
| Alteromonadaceae | 37 | -2.10 | 0.65 | -3.2 | 2 | 3.3e-03 | 1.2e-02 |
| Bdellovibrionaceae | 22 | -1.80 | 0.44 | -4.0 | 2 | 3.8e-04 | 1.9e-03 |
| Caedibacteraceae | 19 | 1.20 | 0.48 | 2.5 | 2 | 1.9e-02 | 4.1e-02 |
| Chitinophagaceae | 220 | 2.80 | 0.53 | 5.2 | 2 | 1.4e-05 | 1.2e-04 |
| Chthoniobacteraceae | 45 | -3.20 | 0.49 | -6.6 | 2 | 4.0e-07 | 6.9e-06 |
| Desulfocapsaceae | 5 | 2.40 | 0.77 | 3.1 | 2 | 4.3e-03 | 1.4e-02 |
| Elsteraceae | 7 | -1.50 | 0.58 | -2.5 | 2 | 1.8e-02 | 4.0e-02 |
| Fimbriimonadaceae | 11 | 3.30 | 0.82 | 4.0 | 2 | 4.0e-04 | 1.9e-03 |
| Flavobacteriaceae | 13000 | 1.80 | 0.53 | 3.4 | 2 | 2.3e-03 | 8.6e-03 |
| Halomonadaceae | 8 | -0.78 | 0.32 | -2.4 | 2 | 2.3e-02 | 4.8e-02 |
| Hydrogenophilaceae | 52 | 1.90 | 0.55 | 3.5 | 2 | 1.4e-03 | 5.7e-03 |
| Hymenobacteraceae | 19 | 1.80 | 0.68 | 2.6 | 2 | 1.6e-02 | 3.5e-02 |
| Hyphomonadaceae | 270 | 1.10 | 0.46 | 2.4 | 2 | 2.1e-02 | 4.3e-02 |
| **Ilumatobacteraceae** | **74** | **-2.70** | **0.42** | **-6.5** | **2** | **5.0e-07** | **6.9e-06** |
| Isosphaeraceae | 28 | 1.20 | 0.47 | 2.6 | 2 | 1.4e-02 | 3.2e-02 |
| Kordiimonadaceae | 12 | 1.30 | 0.49 | 2.7 | 2 | 1.2e-02 | 3.0e-02 |
| Lactobacillaceae | 20 | -1.90 | 0.45 | -4.2 | 2 | 2.7e-04 | 1.5e-03 |
| Methyloligellaceae | 39 | -1.60 | 0.44 | -3.7 | 2 | 8.9e-04 | 3.9e-03 |
| Micavibrionaceae | 82 | -3.50 | 0.31 | -11.0 | 2 | 0.0e+00 | 0.0e+00 |
| **Microbacteriaceae** | **15000** | **-3.40** | **0.40** | **-8.5** | **2** | **0.0e+00** | **1.0e-07** |
| **Microtrichaceae** | **64** | **-2.80** | **0.55** | **-5.1** | **2** | **1.9e-05** | **1.5e-04** |
| Moraxellaceae | 1000 | 1.30 | 0.50 | 2.7 | 2 | 1.3e-02 | 3.2e-02 |
| **Mycobacteriaceae** | **370** | **-1.10** | **0.39** | **-2.8** | **2** | **9.2e-03** | **2.4e-02** |
| NA | 1000 | -1.50 | 0.43 | -3.5 | 2 | 1.5e-03 | 5.9e-03 |
| Nannocystaceae | 48 | -2.20 | 0.74 | -3.0 | 2 | 5.6e-03 | 1.7e-02 |
| **Neisseriaceae** | **240** | **2.10** | **0.66** | **3.1** | **2** | **3.9e-03** | **1.3e-02** |
| Nitrosomonadaceae | 620 | 4.00 | 1.30 | 3.2 | 2 | 3.6e-03 | 1.3e-02 |
| Paracaedibacteraceae | 200 | -3.50 | 0.95 | -3.7 | 2 | 9.4e-04 | 4.0e-03 |
| Parachlamydiaceae | 160 | 1.70 | 0.41 | 4.1 | 2 | 3.2e-04 | 1.7e-03 |
| Pedosphaeraceae | 13 | -1.30 | 0.44 | -2.9 | 2 | 7.4e-03 | 2.1e-02 |
| **Pseudomonadaceae** | **570** | **-2.30** | **0.85** | **-2.7** | **2** | **1.2e-02** | **3.1e-02** |
| Pseudonocardiaceae | 5 | 0.92 | 0.31 | 3.0 | 2 | 5.6e-03 | 1.7e-02 |
| Reyranellaceae | 4000 | 1.90 | 0.34 | 5.4 | 2 | 8.7e-06 | 8.1e-05 |
| **Rhizobiaceae** | **360** | **-3.70** | **0.33** | **-11.0** | **2** | **0.0e+00** | **0.0e+00** |
| **Rhizobiales Incertae Sedis** | **5600** | **-6.50** | **0.99** | **-6.6** | **2** | **4.0e-07** | **6.9e-06** |
| **Rhodanobacteraceae** | **50** | **-3.00** | **0.64** | **-4.6** | **2** | **7.4e-05** | **5.4e-04** |
| Rickettsiaceae | 54 | -2.10 | 0.82 | -2.6 | 2 | 1.5e-02 | 3.4e-02 |
| Rubinisphaeraceae | 14 | -1.40 | 0.50 | -2.8 | 2 | 9.1e-03 | 2.4e-02 |
| Sandaracinaceae | 1 | 4.80 | 0.75 | 6.4 | 2 | 7.0e-07 | 7.7e-06 |
| Saprospiraceae | 62 | -3.00 | 0.78 | -3.9 | 2 | 6.2e-04 | 2.9e-03 |
| Simkaniaceae | 59 | 3.40 | 0.54 | 6.4 | 2 | 6.0e-07 | 7.7e-06 |
| **Solirubrobacteraceae** | **280** | **-2.00** | **0.79** | **-2.5** | **2** | **1.9e-02** | **4.0e-02** |
| Sphingomonadaceae | 3400 | 1.40 | 0.47 | 3.0 | 2 | 5.7e-03 | 1.7e-02 |
| Spirosomaceae | 700 | 2.60 | 0.64 | 4.2 | 2 | 2.7e-04 | 1.5e-03 |
| Sulfuricellaceae | 4 | 2.70 | 0.65 | 4.2 | 2 | 2.6e-04 | 1.5e-03 |
| **Unknown Family** | **1200** | **-4.20** | **0.92** | **-4.6** | **2** | **9.0e-05** | **6.1e-04** |
| Verrucomicrobiaceae | 86 | 4.10 | 0.54 | 7.5 | 2 | 0.0e+00 | 1.0e-06 |
| ***Origin (LP vs HP)*** | | | | | | | |
| Rhodobacteraceae | 5100 | 1.70 | 0.30 | 5.6 | 2 | 5.6e-06 | 5.7e-04 |

### Supplementary material 13. Summary of the results of the linear models for differential abundance analysis (LINDA) testing the effects of metal contamination (NC vs. C), immune challenge (PBS vs. AMIX), and the fish origin (NC vs. C) on water (within fish tanks) microbiome composition at the phylum level. Analysis was performed on original log-transformed data. Only the taxa that have been significantly (adjusted p < 0.05 and |log2FoldChange| > 0.5) differentially abundant among treatments are displayed. Taxa for which relative abundance changes according to contamination treatment in both water and gut are highlighted in bold.

| ***Differential abundance of Taxa at the Phylum level – LINDA results for water*** | | | | | | | |
| --- | --- | --- | --- | --- | --- | --- | --- |
|  | baseMean | log2FoldChange | lfcSE | stat | df | pvalue | padj |
| ***Contamination (NC vs. C)*** | | | | | | | |
| **Actinobacteriota** | **16000** | **-3.00** | **0.31** | **-9.6** | **2** | **0.0e+00** | **0.0e+00** |
| Bacteroidota | 12000 | 1.10 | 0.43 | 2.5 | 2 | 1.7e-02 | 3.7e-02 |
| Bdellovibrionota | 16 | -2.80 | 0.39 | -7.1 | 2 | 1.0e-07 | 4.0e-07 |
| Firmicutes | 400 | -3.00 | 0.45 | -6.5 | 2 | 5.0e-07 | 1.7e-06 |
| NA | 140 | -5.80 | 0.35 | -17.0 | 2 | 0.0e+00 | 0.0e+00 |
| Patescibacteria | 19 | -2.10 | 0.53 | -4.0 | 2 | 3.8e-04 | 9.4e-04 |
| **Proteobacteria** | **71000** | **-0.63** | **0.27** | **-2.4** | **2** | **2.5e-02** | **4.7e-02** |
| Verrucomicrobiota | 310 | 1.60 | 0.32 | 4.9 | 2 | 3.5e-05 | 1.1e-04 |

### Supplementary material 14. Summary of the results of the linear models for differential abundance analysis (LINDA) testing the effects of metal contamination (NC vs. C), immune challenge (PBS vs. AMIX), and the fish origin (NC vs. C) on functions inferred from fish gut microbiome sequences (MOTUs). Analysis was performed on original log-transformed data. Only the functions that have been significantly (adjusted p < 0.05 and |log2FoldChange| > 0.5) differentially abundant among treatments are displayed.

| ***Differential abundance of inferred functions - LINDA Results*** | | | | | | | |
| --- | --- | --- | --- | --- | --- | --- | --- |
|  | baseMean | log2FoldChange | lfcSE | stat | df | pvalue | padj |
| ***Contamination (NC vs. C)*** | | | | | | | |
| Anaerobic Respiration | 6.9 | 0.81 | 0.28 | 2.9 | 290 | 3.5e-03 | 3.8e-02 |
| C1 Compounds | 470.0 | -0.62 | 0.20 | -3.1 | 66 | 3.0e-03 | 3.8e-02 |
| Cell Structure Biosynthesis | 2100.0 | -0.69 | 0.22 | -3.1 | 63 | 2.6e-03 | 3.8e-02 |
| Demethylmenaquinone Biosynthesis | 1900.0 | -0.62 | 0.18 | -3.4 | 62 | 1.1e-03 | 1.9e-02 |
| Secondary Metabolite Biosynthesis | 1100.0 | -0.75 | 0.21 | -3.5 | 59 | 8.1e-04 | 1.8e-02 |
| Sugar Alcohols Degradation | 27.0 | 0.59 | 0.20 | 2.9 | 68 | 5.3e-03 | 4.7e-02 |
| Superpathway Of Chorismate Metabolism | 1300.0 | -0.84 | 0.19 | -4.4 | 60 | 4.4e-05 | 2.0e-03 |
| ***Size (cm)*** | | | | | | | |
| Vitamin Degradation | 3.2 | 0.54 | 0.11 | 5.0 | 300 | 9.0e-07 | 7.8e-05 |

### Supplementary material 15. Comprehensive summary of the specific pathways affected by the experimental metal contamination. Pathway classes reported as "Lower-abundant" have shown a reduction in their relative abundance under experimental exposure to metal contamination (C) compared to noncontaminated treatment (NC), suggesting a possible inhibitory effect of metal contamination on these metabolic functions. Conversely, pathway classes labeled "Over-abundant" have increased relative abundance, indicating a potential stimulation of the function under metal contamination exposure. Each entry lists the pathway class alongside a brief description and its parent class to illustrate broader biochemical categories.

| Effect of exposure to metal contamination | Pathway class | Description | parent class |
| --- | --- | --- | --- |
| Lower-abundant | C1 Compounds | This class involves the utilization of C1 compound pathways such as CO2 fixation, carbon monoxide utilization, formaldehyde utilization, and methanol utilization. | Degradation/Utilization/Assimilation |
|  | Demethylmenaquinone Biosynthesis | This class contains pathways for the biosynthesis of demethylmenaquinones - a low-molecular-weight lipophilic component of the cytoplasmic membrane of bacteria mediating electron transfer between hydrogenases and cytochrome. | Quinol and Quinone Biosynthesis |
|  | Secondary Metabolite Biosynthesis | This class contains pathways involved in the biosynthesis of secondary metabolites, organic compounds that are not directly involved in organisms' growth, development, and reproduction. | Biosynthesis |
|  | Superpathway Of Chorismate Metabolism | This class contains pathways involved in the metabolism and biosynthesis of chorismate. More especially, chorismate is the principal common precursor of the aromatic amino acids tryptophan, tyrosine, and phenylalanine, as well as the essential compounds such as tetrahydrofolate, ubiquinone-8, menaquinone-8, and enterobactin (enterochelin). | Biosynthesis |
|  | Cell Structure Biosynthesis | This class contains pathways involved in the biosynthesis of cellular organelles, including cell wall components and substances associated with them. | Biosynthesis |
| Over-abundant | Anaerobic Respiration | This class contains pathways involved in the enzymatic release of energy from inorganic and organic compounds (especially carbohydrates and fats) by using compounds other than oxygen (e.g., nitrate, sulfate) as the terminal electron acceptor. | Respiration |
|  | Sugar Alcohols Degradation | This class contains pathways involved in using various sugars & alcohols as carbon and energy sources. | Alcohol Degradation & Carbohydrate Degradation |

### References

1.	Schliep KP. phangorn: phylogenetic analysis in R. *Bioinformatics* 2011; **27**: 592–593.

2.	Wright ES. Using DECIPHER v2.0 to Analyze Big Biological Sequence Data in R. *The R Journal* 2016; **8**: 352–359.

3.	Nearing JT, Douglas GM, Hayes MG, MacDonald J, Desai DK, Allward N, et al. Microbiome differential abundance methods produce different results across 38 datasets. *Nat Commun* 2022; **13**: 342.

4.	Cappellato M, Baruzzo G, Di Camillo B. Investigating differential abundance methods in microbiome data: A benchmark study. *PLoS Comput Biol* 2022; **18**: e1010467.

5.	Schloss PD. Waste not, want not: Revisiting the analysis that called into question the practice of rarefaction. 2023. Microbiology.

6.	Gloor GB, Macklaim JM, Pawlowsky-Glahn V, Egozcue JJ. Microbiome Datasets Are Compositional: And This Is Not Optional. *Frontiers in Microbiology* 2017; **8**.

7.	McMurdie PJ, Holmes S. Waste Not, Want Not: Why Rarefying Microbiome Data Is Inadmissible. *PLOS Computational Biology* 2014; **10**: e1003531.
